# Batch effects and the effective design of single-cell gene expression studies

**DOI:** 10.1101/062919

**Authors:** Po-Yuan Tung, John D. Blischak, Chiaowen Joyce Hsiao, David A. Knowles, Jonathan E. Burnett, Jonathan K. Pritchard, Yoav Gilad

## Abstract

Single cell RNA sequencing (scRNA-seq) can be used to characterize variation in gene expression levels at high resolution. However, the sources of experimental noise in scRNA-seq are not yet well understood. We investigated the technical variation associated with sample processing using the single cell Fluidigm C1 platform. To do so, we processed three C1 replicates from three human induced pluripotent stem cell (iPSC) lines. We added unique molecular identifiers (UMIs) to all samples, to account for amplification bias. We found that the major source of variation in the gene expression data was driven by genotype, but we also observed substantial variation between the technical replicates. We observed that the conversion of reads to molecules using the UMIs was impacted by both biological and technical variation, indicating that UMI counts are not an unbiased estimator of gene expression levels. Based on our results, we suggest a framework for effective scRNA-seq studies.

## Introduction

Single-cell genomic technologies can be used to study the regulation of gene expression at unprecedented resolution [1,2]. Using single-cell gene expression data, we can begin to effectively characterize and classify individual cell types and cell states, develop a better understanding of gene regulatory threshold effects in response to treatments or stress, and address a large number of outstanding questions that pertain to the regulation of noise and robustness of gene expression programs. Indeed, single cell gene expression data have already been used to study and provide unique insight into a wide range of research topics, including differentiation and tissue development [3–5], the innate immune response [6,7], and pharmacogenomics [8,9].

Yet, there are a number of outstanding challenges that arose in parallel with the application of single cell technology [10]. A fundamental difficulty, for instance, is the presence of inevitable technical variability introduced during sample processing steps, including but not limited to the conditions of mRNA capture from a single cell, amplification bias, sequencing depth, and variation in pipetting accuracy. These (and other sources of error) may not be unique to single cell technologies, but in the context of studies where each sample corresponds to a single cell, and is thus processed as a single unrepeatable batch, these technical considerations make the analysis of biological variability across single cells particularly challenging.

To better account for technical variability in scRNA-seq experiments, it has become common to add spike-in RNA standards of known abundance to the endogenous samples [11,12]. The most commonly used spike-in was developed by the External RNA Controls Consortium (ERCC) [13]; comprising of a set of 96 RNA controls of varying length and GC content. A number of single cell studies focusing on analyzing technical variability based on ERCC spike-in controls have been reported [11,12,14,15]. However, one principle problem with spike-ins is that they do not ‘experience’ all processing steps that the endogenous sample is subjected to. For that reason, it is unknown to what extent the spike-ins can faithfully reflect the error that is being accumulated during the entire sample processing procedure, either within or across batches. In particular, amplification bias, which is assumed to be gene-specific, cannot be addressed by spike-in normalization approaches.

To address challenges related to the efficiency and uniformity with which mRNA molecules are amplified and sequenced in single cells, unique molecule identifiers (UMIs) were introduced to single cell sample processing [16–19]. The rationale is that by counting molecules rather than the number of amplified sequencing reads, one can account for biases related to amplification, and obtain more accurate estimates of gene expression levels [7,12,20]. It is assumed that most sources of variation in single cell gene expression studies can be accounted for by using the combination of UMIs and a spike-in based standardization [15,20]. Nevertheless, though molecule counts, as opposed to sequencing read counts, are associated with substantially reduced levels of technical variability, a non-negligible proportion of experimental error remains unexplained.

There are a few common platforms in use for scRNA-seq. The automated C1 microfluidic platform (Fluidigm), while more expensive per sample, has been shown to confer several advantages over platforms that make use of droplets to capture single cells [3,21]. In particular, smaller samples can be processed using the C1 (when cell numbers are limiting), and the C1 capture efficiency of genes (and RNA molecules) is markedly higher. Notably, in the context of this study, the C1 system also allows for direct confirmation of single cell capture events, in contrast to most other microfluidic-based approaches [3,22]. One of the biggest limitations of using the C1 system, however, is that single cell capture and preparation from different conditions are fully independent [23]. Consequently, multiple replicates of C1 collections from the same biological condition are necessary to facilitate estimation of technical variability even with the presence of ERCC spike-in controls [10]. To our knowledge, to date, no study has been purposely conducted to assess the technical variability across batches on the C1 platform.

To address this gap, we collected scRNA-seq data from induced pluripotent stem cell (iPSC) lines of three Yoruba individuals (abbreviation: YRI) using C1 microfluidic plates. Specifically, we performed three independent C1 collections per each individual to disentangle batch effects from the biological covariate of interest, which, in this case, is the difference between individuals. Both ERCC spike-in controls and UMIs were included in our sample processing. With these data, we were able to elucidate technical variability both within and between C1 batches and thus provide a deep characterization of cell-to-cell variation in gene expression levels across individuals.

## Results

### Study design and quality control

We collected single cell RNA-seq (scRNA-seq) data from three YRI iPSC lines using the Fluidigm C1 microfluidic system followed by sequencing. We added ERCC spike-in controls to each sample, and used 5-bp random sequence UMIs to allow for the direct quantification of mRNA molecule numbers. For each of the YRI lines, we performed three independent C1 collections; each replicate was accompanied by processing of a matching bulk sample using the same reagents. This study design (Figure 1A and Table S1A) allows us to estimate error and variability associated with the technical processing of the samples, independently from the biological variation across single cells of different individuals. We were also able to estimate how well scRNA-seq data can recapitulate the RNA-seq results from population bulk samples.

In what follows, we describe data as originating from different samples when we refer to data from distinct wells of each C1 collection. Generally, each sample corresponds to a single cell. In turn, we describe data as originating from different replicates when we refer to all samples from a given C1 collection, and from different individuals when we refer to data from all samples and replicates of a given genetically distinct iPSC line.

We obtained an average of 6.3 +/− 2.1 million sequencing reads per sample (range 0.411.2 million reads). We processed the sequencing reads using a standard alignment approach (see Methods) and performed multiple quality control analyses. As a first step, we estimated the proportion of ERCC spike-in reads from each sample. We found that, across samples, sequencing reads from practically all samples of the second replicate of individual NA19098 included unusually high ERCC content compared to all other samples and replicates (Figure S1A and S1B). We concluded that a pipetting error led to excess ERCC content in this replicate and we excluded the data from all samples of this replicate in subsequent analyses. With the exception of the excluded samples, data from all other replicates seem to have similar global properties (using general metrics; Figure 1C-E and Figure S1C-F).

We next examined the assumption that data from each sample correspond to data from a single cell. After the cell sorting was complete, but before the processing of the samples, we performed visual inspection of the C1 microfluidic plates. Based on that visual inspection, we flagged 21 samples that did not contain any cell, and 54 samples that contained more than one cell (across all batches). Visual inspection of the C1 microfluidic plate is an important quality control step, but it is not infallible. We therefore filtered data from the remaining samples based on the number of total mapped reads, the percentage of unmapped reads, the percentage of ERCC spike-in reads, and the number of genes detected (Figure 1B-E). We chose data-driven inclusion cutoffs for each metric, based on the 95th percentile of the respective distributions for the 21 libraries that were amplified from samples that did not include a cell based on visual inspection (Figure S1C-F). Using this approach, we identified and removed data from 15 additional samples that were classified as originating from a single cell based on visual inspection, but whose data were more consistent with a multiple-cell origin based on the number of total molecules, the concentration of cDNA amplicons, and the read-to-molecule conversion efficiency (defined as the number of total molecules divided by the number of total reads; Figure S2). At the conclusion of these quality control analyses and exclusion steps, we retained data from 564 high quality samples, which correspond, with reasonable confidence, to 564 single cells, across eight replicates from three individuals (Table S2).

Our final quality check focused on the different properties of sequencing read and molecule count data. We considered data from the 564 high quality samples and compared gene specific counts of sequencing read and molecules. We found that while gene-specific reads and molecule counts are exceptionally highly correlated when we considered the ERCC spike-in data (r = 0.99; Figure 1F), these counts are somewhat less correlated when data from the endogenous genes are considered (r = 0.92). Moreover, the gene-specific read and molecule counts correlation is noticeably lower for genes that are expressed at lower levels (Figure 1F). These observations concur with previous studies [12,20] as they underscore the importance of using UMIs in single cell gene expression studies.

**Figure 1.**
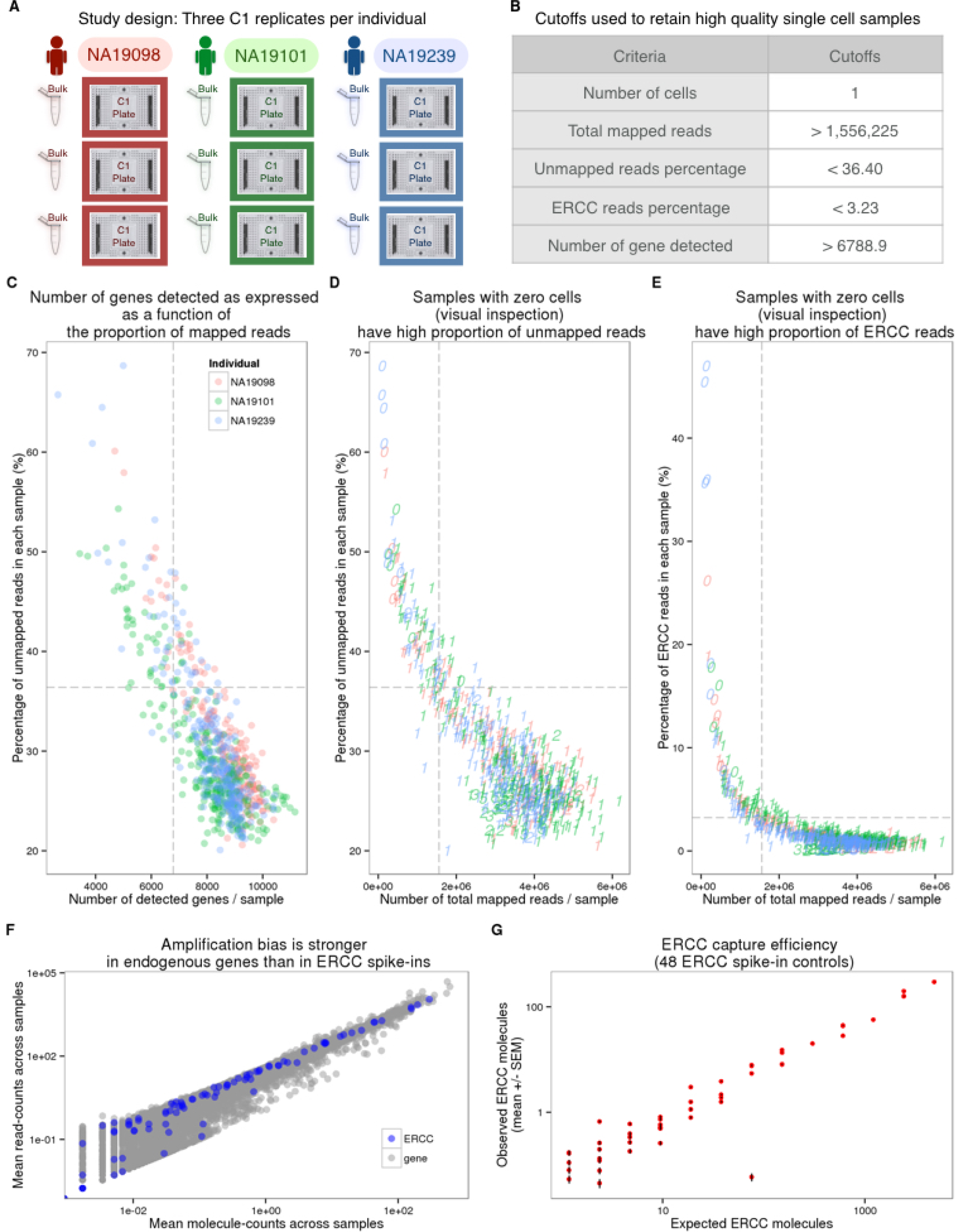
Experimental design and quality control of scRNA-seq. (A) Three C1 96 well-integrated fluidic circuit (IFC) replicates were collected from each of the three Yoruba individuals. A bulk sample was included in each batch. (B) Summary of the cutoffs used to remove data from low quality cells that might be ruptured or dead (See Figure S1 for details). (C-E) To assess the quality of the scRNA-seq data, the capture efficiency of cells and the faithfulness of mRNA fraction amplification were determined based on the proportion of unmapped reads, the number of detected genes, the numbers of total mapped reads, and the proportion of ERCC spike-in reads across cells. The dash lines indicate the cutoffs summarized in panel (B). The three colors represent the three individuals (NA19098 in red, NA19101 in green, and NA19239 in blue), and the numbers indicate the cell numbers observed in each capture site on C1 plate. (F) Scatterplots in log scale showing the mean read counts and the mean molecule counts of each endogenous gene (grey) and ERCC spike-ins (blue) from the 564 high quality single cell samples before removal of genes with low expression. (G) mRNA capture efficiency shown as observed molecule count versus number of molecules added to each sample, only including the 48 ERCC spike-in controls remaining after removal of genes with low abundance. Each red dot represents the mean +/− SEM of an ERCC spike-in across the 564 high quality single cell samples.

We proceeded by investigating the effect of sequencing depth and the number of single cells collected on multiple properties of the data. To this end, we repeatedly subsampled single cells and sequencing reads to assess the correlation of the single cell gene expression estimates to the bulk samples, the number of genes detected, and the correlation of the cell-to-cell gene expression variance estimates between the reduced subsampled data and the full single cell gene expression data set (Figure 2). We observed quickly diminishing improvement in all three properties with increasing sequencing depth and the number of sampled cells, especially for highly expressed genes. For example, a per cell sequencing depth of 1.5 million reads (which corresponds to ~50,000 molecules) from each of 75 single cells was sufficient for effectively quantifying even the lower 50% of expressed genes. At this level of subsampling for individual NA19239, we were able to detect a mean of 6068 genes out of 6097 genes expressed in the bulk samples (the bottom 50%; Figure 2B); the estimated single cell expression levels of these genes (summed across all cells) correlated with the bulk sample gene expression levels with a mean Pearson coefficient of 0.8 (Figure 2A), and the estimated cell-to-cell variation in gene expression levels was correlated with the variation estimated from the full data set with a mean Pearson coefficient of 0.95 (Figure 2C).

**Figure 2.**
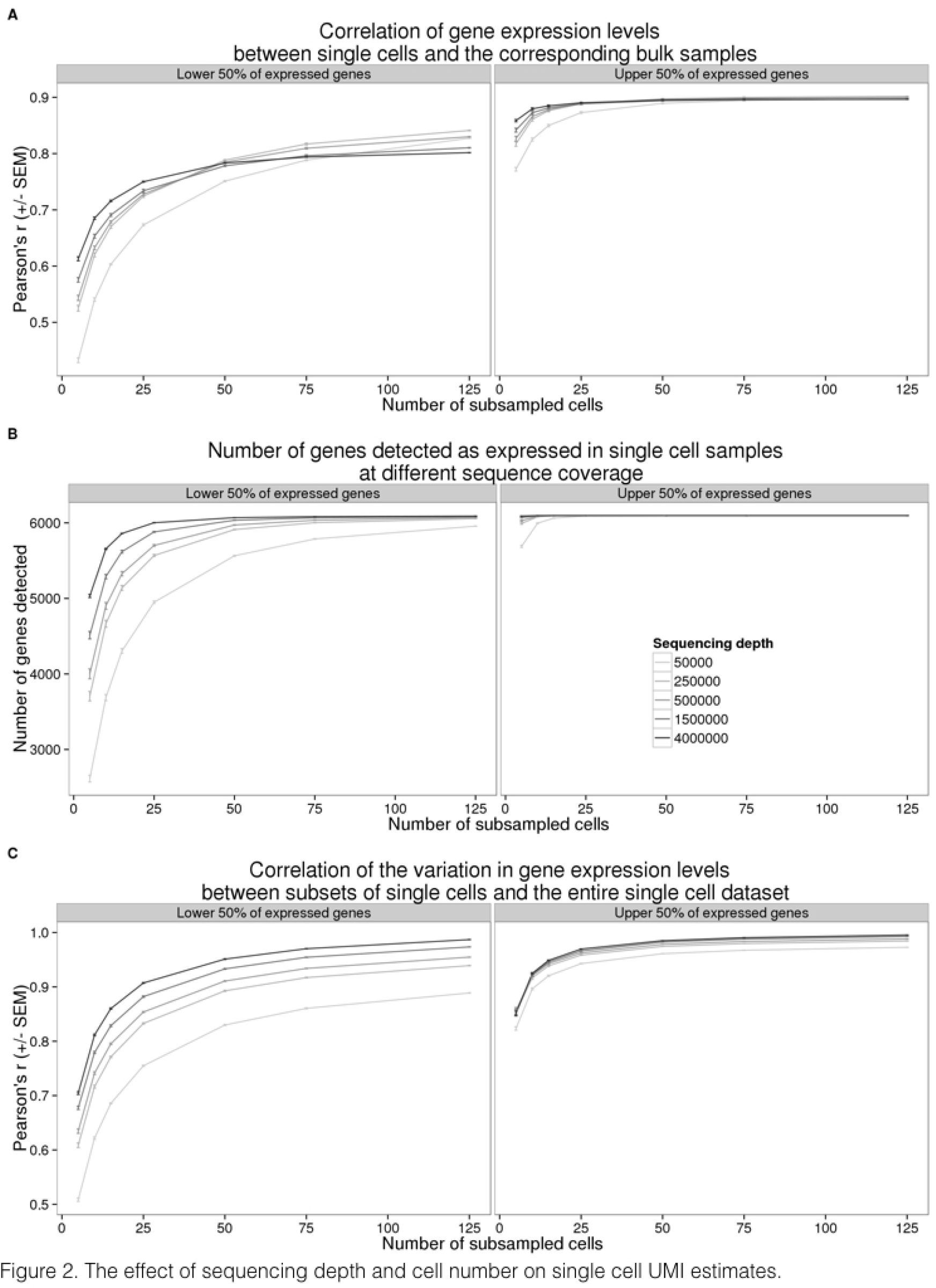
The effect of sequencing depth and cell number on single cell UMI estimates. Sequencing reads from the entire data set were subsampled to the indicated sequencing depth and cell number, and subsequently converted to molecules using the UMIs. Each point represents the mean +/− SEM of 10 random draws of the indicated cell number. The left panel displays the results for 6,097 (50% of detected) genes with lower expression levels and the right panel the results for 6,097 genes with higher expression levels. (A) Pearson correlation of aggregated gene expression level estimates from single cells compared to the bulk sequencing samples. (B) Total number of genes detected with at least one molecule in at least one of the single cells. (C) Pearson correlation of cell-to-cell gene expression variance estimates from subsets of single cells compared to the full single cell data set.

### Batch effects associated with UMI-based single cell data

In the context of the C1 platform, typical study designs make use of a single C1 plate (batch/replicate) per biological condition. In that case, it is impossible to distinguish between biological and technical effects associated with the independent capturing and sequencing of each C1 replicate. We designed our study with multiple technical replicates per biological condition (individual) in order to directly and explicitly estimate the batch effect associated with independent C1 preparations (Figure 1A).

As a first step in exploring batch effects, we examined the gene expression profiles across all single cells that passed our quality checks (as reported above) using raw molecule counts (without standardization). Using principal component analysis (PCA) for visualization, we observed-as expected-that the major source of variation in data from single cells is the individual origin of the sample (Figure 4A). Specifically, we found that the proportion of variance due to individual was larger (median: 8%) than variance due to C1 batch (median: 4%; Kruskal-Wallis test; *P* < 0.001, Figure S3A; see Methods for details of the variance component analysis). Yet, variation due to C1 batch is also substantial - data from single cell samples within a batch are more correlated than that from single cells from the same individual but different batches (Kruskal-Wallis test; *P* < 0.001).

Could we account for the observed batch effects using the ERCC spike-in controls? In theory, if the total ERCC molecule-counts are affected only by technical variability, the spike-ins could be used to correct for batch effects even in a study design that entirely confounds biological samples with C1 preparations. To examine this, we first considered the relationship between total ERCC molecule-counts and total endogenous molecule-counts per sample. If only technical variability affects ERCC molecule-counts, we expect the technical variation in the spike-ins (namely, variation between C1 batches) to be consistent, regardless of the individual assignment. Indeed, we observed that total ERCC molecule-counts are significantly different between C1 batches (F-test; *P* < 0.001). However, total ERCC molecule-counts are also quite different across individuals, when variation between batches is taken into account (LRT; *P* = 0. 08; Figure 3A). This observation suggests that both technical and biological variation affect total ERCC molecule-counts. In addition, while we observed a positive relationship between total ERCC molecule-counts and total endogenous molecule-counts per sample, this correlation pattern differed across C1 batches and across individuals (F-test; *P* < 0.001; Figure 3B).

To more carefully examine the technical and biological variation of ERCC spike-in controls, we assessed the ERCC per-gene expression profile. We observed that the ERCC gene expression data from samples of the same batch were more correlated than data from samples across batches (Kruskal-Wallis test; Chi-squared *P* < 0.001). However, the proportion of variance explained by the individual was significantly larger than the variance due to C1 batch (median: 9% vs. 5%, Chi-squared test; *P* < 0.001, Figure S3B), lending further support to the notion that biological variation affects the ERCC spike in data. Based on these analyses, we concluded that ERCC spike-in controls cannot be used to effectively account for the batch effect associated with independent C1 preparations.

We explored potential reasons for the observed batch effects, and in particular, the difference in ERCC counts across batches and individuals. We focused on the read-to-molecule conversion rates, i.e. the rates at which sequencing reads are converted to molecule counts based on the UMI sequences. We defined read-to-molecule conversion efficiency as the total molecule-counts divided by the total reads-counts in each sample, considering separately the reads/molecules that correspond to endogenous genes or ERCC spike-ins (Figure 3C and 3D). We observed a significant batch effect in the read-to-molecule conversion efficiency of both ERCC (F-test; *P* < 0.05) and endogenous genes (F-test; *P* < 0.001) across C1 replicates from the same individual. Moreover, the difference in read-to-molecule conversion efficiency across the three individuals was significant not only for endogenous genes (LRT; *P* < 0.01, Figure 3C) but also in the ERCC spike-ins (LRT; *P* < 0.01, Figure 3D). We reason that the difference in read to molecule conversion efficiency across C1 preparations may contribute to the observed batch effect in this platform.

**Figure 3.**
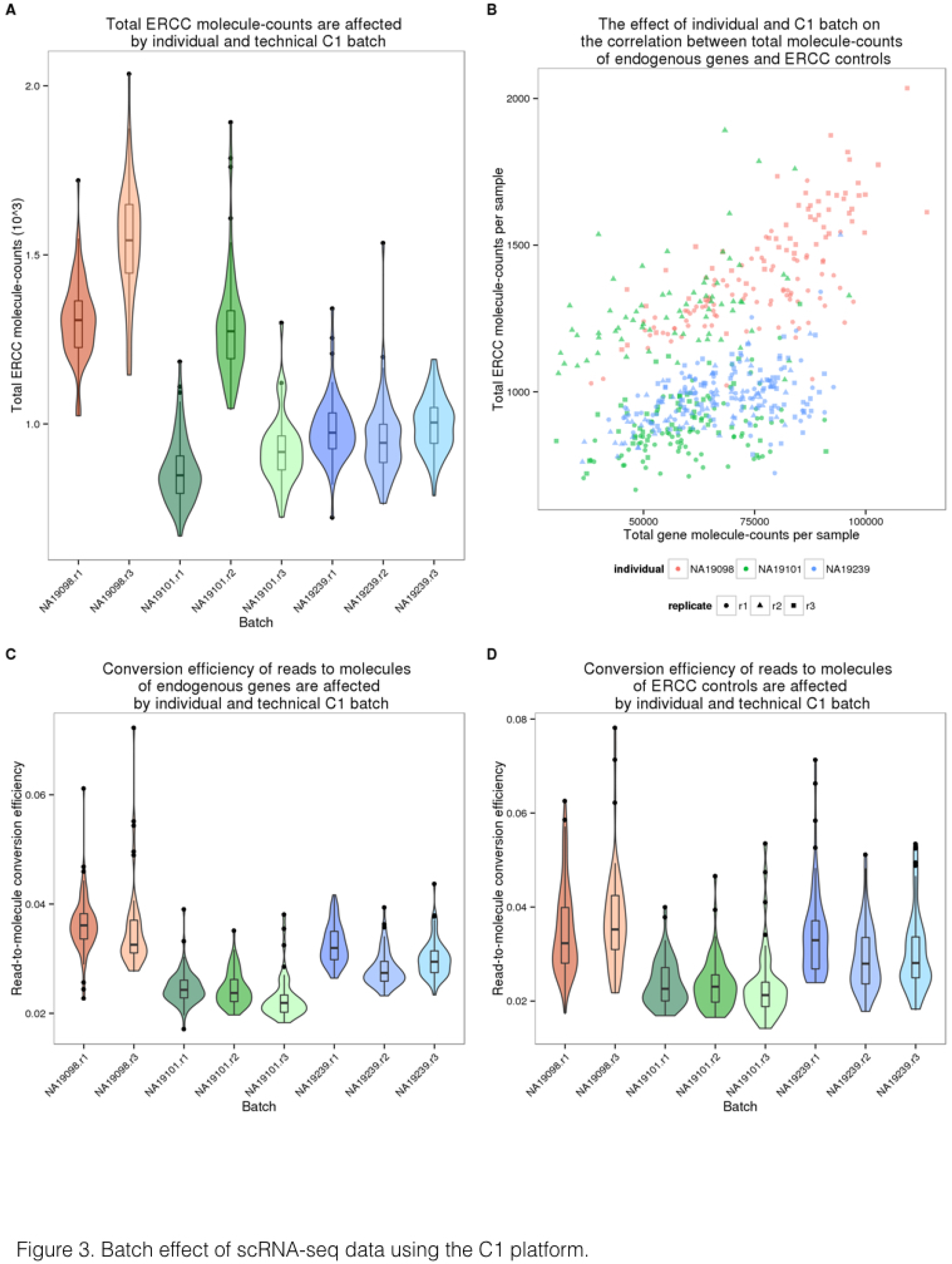
Batch effect of scRNA-seq data using the C1 platform. (A) Violin plots of the number of total ERCC spike-in molecule-counts in single cell samples per C1 replicate. (B) Scatterplot of the total ERCC molecule-counts and total gene molecule-counts. The colors represent the three individuals (NA19098 is in red, NA19101 in green, and NA19239 in blue). Data from different C1 replicates is plotted in different shapes. (C and D) Violin plots of the reads to molecule conversion efficiency (total molecule-counts divided by total read-counts per single cells) by C1 replicate. The endogenous genes and the ERCC spike-ins are shown separately in (C) and (D), respectively. There is significant difference across individuals of both endogenous genes (*P* < 0.001) and ERCC spike-ins (*P* < 0.05). The differences across C1 replicates per individual of endogenous genes and ERCC spike-ins were also evaluated (both *P* < 0.01).

### Measuring regulatory noise in single-cell gene expression data

Our analysis indicated that there is a considerable batch effect in the single cell gene expression data collected from the C1 platform. We thus sought an approach that would account for the batch effect and allow us to study biological properties of the single-cell molecule count-based estimates of gene expression levels, albeit in a small sample of just three individuals. As a first step, we adjusted the raw molecule counts by using a Poisson approximation to account for the random use of identical UMI sequences in molecules from highly expressed genes (this was previously termed a correction for the UMI ‘collision probability’ [17]). We then excluded data from genes whose inferred molecule count exceeded 1,024 (the theoretical number of UMI sequences) – this step resulted in the exclusion of data from 6 mitochondrial genes.

We next incorporated a standardization step by computing log transformed counts-per-million (cpm) to remove the effect of different sequencing depths, as is the common practice for the analysis of bulk RNA-seq data (Figure 4A and 4B). We used a Poisson generalized linear model to normalize the endogenous molecule log_2_ cpm values by the observed molecule counts of ERCC spike-ins across samples. While we do not expect this step to account for the batch effect (as discussed above), we reasoned that the spike-ins allow us to account for a subset of technical differences between samples, for example, those that arise from differences in RNA concentration (Figure 4C).

Finally, to account for the technical batch effect, we modeled between-sample correlations in gene expression within C1 replicates (see Methods). Our approach is similar in principle to limma, which was initially developed for adjusting within-replicate correlations in microarray data [24]. We assume that samples within each C1 replicate share a component of technical variation, which is independent of biological variation across individuals. We fit a linear mixed model for each gene, which includes a fixed effect for individual and a random effect for batch. The batch effect is specific to each C1 replicate, and is independent of biological variation across individuals. We use this approach to estimate and remove the batch effect associated with different C1 preparations (Figure 4D).

**Figure 4.**
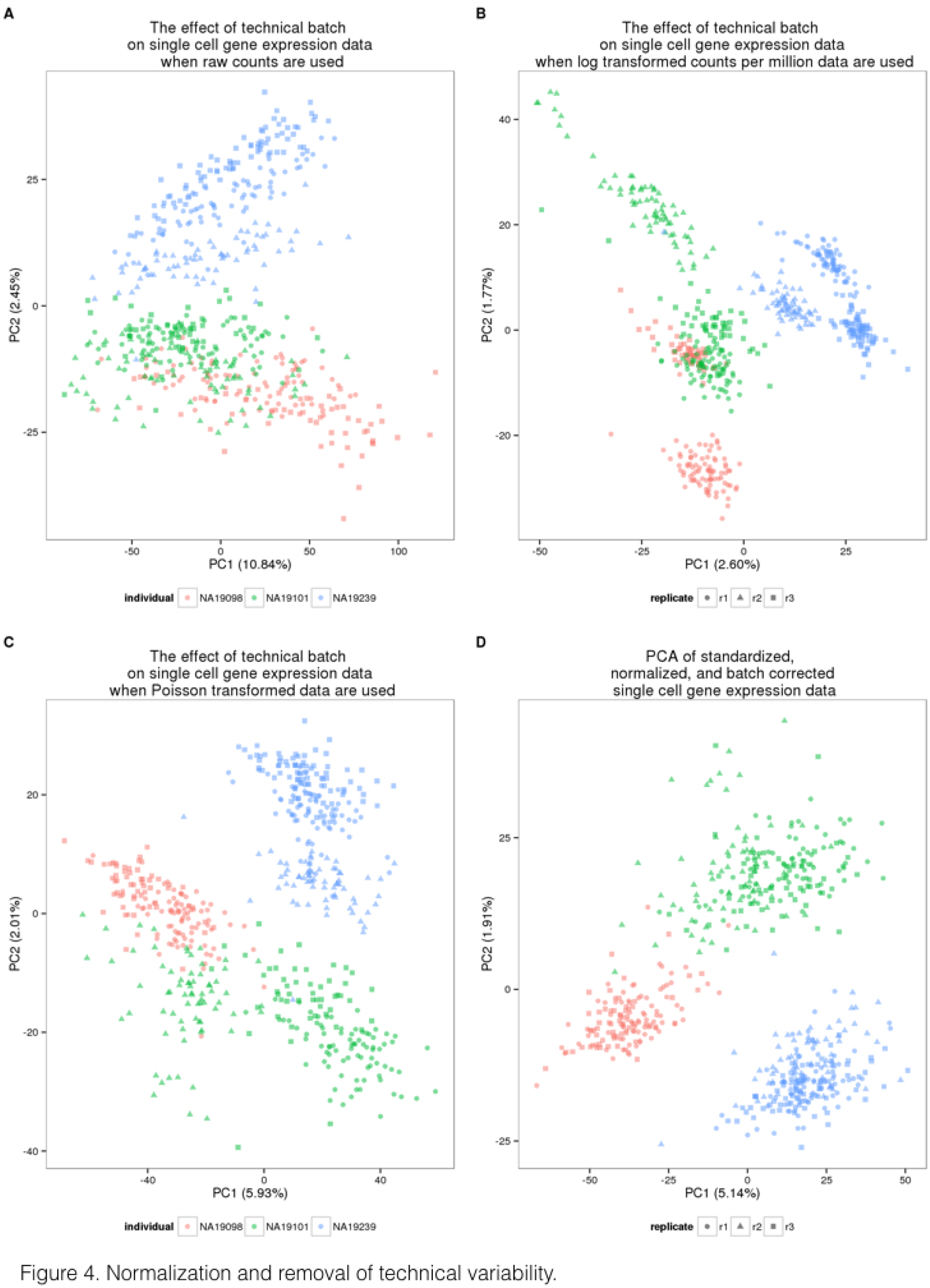
Normalization and removal of technical variability. Principal component (PC) 1 versus PC2 of the (A) raw molecule counts, (B) log_2_ counts per million (cpm), (C) Poisson transformed expression levels (accounting for technical variability modeled by the ERCC spike-ins), and (D) batch-corrected expression levels. The colors represent the three individuals (NA19098 in red, NA19101 in green, and NA19239 in blue). Data from different C1 replicates is plotted in different shapes.

Once we removed the unwanted technical variability, we focused on analyzing biological variation in gene expression between single cells. Our goal was to identify inter-individual differences in the amount of variation in gene expression levels across single cells, or in other words, to identify differences between individuals in the amount of regulatory noise [25]. In this context, regulatory noise is generally defined as the coefficient of variation (CV) of the gene expression levels of single cells [26]. In the following, we used the standardized, normalized, batch-corrected molecule count gene expression data to estimate regulatory noise (Figure 4D). To account for heteroscedasticity from Poisson sampling, we adjusted the CV values by the average gene-specific expression level across cells of the same individual. The adjusted CV is robust both to differences in gene expression levels, as well as to the proportion of gene dropouts in single cells.

To investigate the effects of gene dropouts (the lack of molecule representation of an expressed gene [6,11]) on our estimates of gene expression noise, we considered the association between the proportion of cells in which a given gene is undetected (namely, the gene-specific dropout rate), the average gene expression level, and estimates of gene expression noise. Across all genes, the median gene-specific dropout was 22 percent. We found significant individual differences (LRT; *P* < 10^−5^) in gene-specific dropout rates between individuals in more than 10% (1,214 of 13,058) of expressed endogenous genes. As expected, the expression levels, and the estimated variation in expression levels across cells, are both associated with gene-specific dropout rates (Figure S4A and S4B, respectively). However, importantly, adjusted CVs are not associated with dropout rates (Spearman’s correlation = 0.04; Figure S4C), indicating that adjusted CV measurements are not confounded by the dynamic range of single-cell gene expression levels.

**Figure 5.**
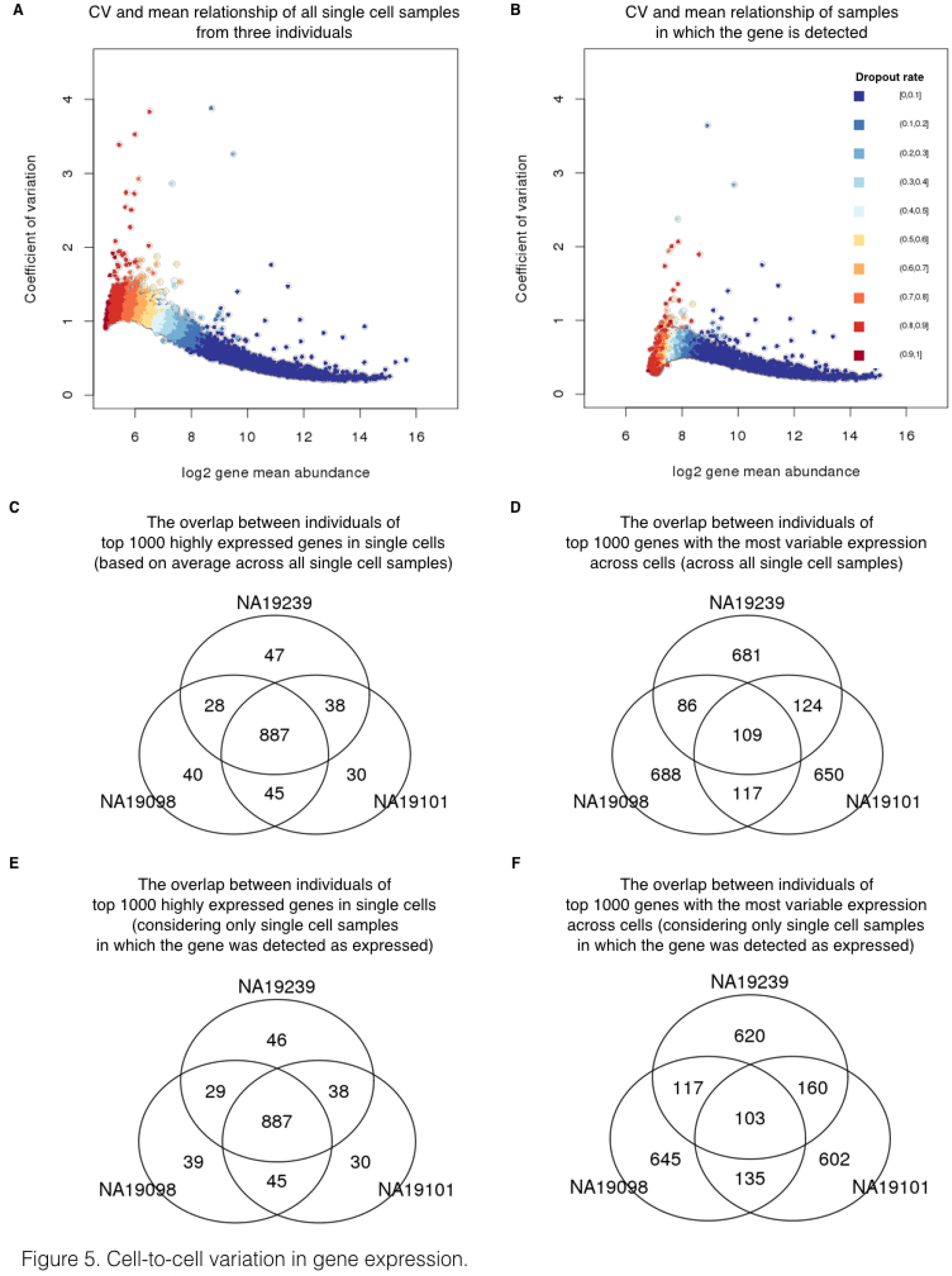
Cell-to-cell variation in gene expression. Adjusted CV plotted against average molecule counts across all cells in (A) and across only the cells in which the gene is expressed (B), including data from all three individuals. Each dot represents a gene, and the color indicates the corresponding gene-specific dropout rate (the proportion of cells in which the gene is undetected). (C and D) Venn diagrams showing the overlaps of top 1000 genes across individuals based on mean expression level in (C) and based on adjusted CV values in (D), considering only the cells in which the gene is expressed. (E and F) Similarly, Venn diagrams showing the overlaps of top 1000 genes across individuals based on mean expression level in (E) and based on adjusted CV values in (F), across all cells.

We thus estimated mean expression levels and regulatory noise (using adjusted CV) for each gene, by either including (Figure 5A) or excluding (Figure 5B) samples in which the gene was not detected/expressed. We first focused on general trends in the data. We ranked genes in each individual by their mean expression level as well as by their estimated level of variation across single cells. When we considered samples in which a gene was expressed, we found that 887 of the 1,000 most highly expressed genes in each individual are common to all three individuals (Figure 5C). In contrast, only 103 of the 1,000 most highly variable (noisy) genes in each individual were common to all three individuals (Figure 5D). We found similar results when we considered data from all single cells, regardless of whether the gene was detected as expressed (Figure 5E and 5F).

Next, we identified genes whose estimated regulatory noise (based on the adjusted CV) is significantly different between individuals. For the purpose of this analysis, we only included data from cells in which the gene was detected as expressed. Based on permutations (Figure S5), we classified the estimates of regulatory noise of 560 genes as significantly different across individuals (empirical *P* < .0001, Figure S6 for examples; Table S3 for gene list). These 560 genes are enriched for genes involved in protein translation, protein disassembly, and various biosynthetic processes (Table S4). Interestingly, among the genes whose regulatory noise estimates differ between individuals, we found two pluripotency genes, *KLF4* and *DPPA2* (Figure S6; Figure S7).

## Discussion

### Study design and sample size for scRNA-seq

Our nested study design allowed us to explicitly estimate technical batch effects associated with single cell sample processing on the C1 platform. We found previously unreported technical sources of variation associated with the C1 sample processing and the use of UMIs, including the property of batch-specific read-to-molecule conversion efficiency. As we used a well-replicated nested study design, we were able to model, estimate, and account for the batch while maintaining individual differences in gene expression levels. We believe that our observations indicate that future studies should avoid confounding C1 batch and individual source of single cell samples. Instead, we recommend a balanced study design consisting of multiple individuals within a C1 plate and multiple C1 replicates (for example, Figure S8A). The origin of each cell can then be identified using the RNA sequencing data. Indeed, using a method originally developed for detecting sample swaps in DNA sequencing experiments [27], we were able to correctly identify the correct YRI individual of origin for all the single cells from the current experiment by comparing the polymorphisms identified using the RNA-seq reads to the known genotypes for all 120 YRI individuals of the International HapMap Project [28] (Figure S8B). The mixed-individual-plate is an attractive study design because it allows one to account for the batch effect without the requirement to explicitly spend additional resources on purely technical replication (because the total number of cells assayed from each individual can be equal to a design in which one individual is being processed in using a single C1 plate).

We also addressed additional study design properties with respect to the desired number of single cells and the desired depth of sequencing (Figure 2). Similar assessments have been previously performed for single cell sequencing with the C1 platform without the use of UMIs [21,29], but no previous study has investigated the effects of these parameters for single cells studies using UMIs. We focused on recapitulating the gene expression levels observed in bulk sequencing experiments, detecting as many genes as possible, and accurately measuring the cell-to-cell variation in gene expression levels. We recommend sequencing at least 75 high quality cells per biological condition with a minimum of 1.5 million raw reads per cell to obtain optimal performance of these three metrics.

### The limitations of the ERCC spike-in controls

The ERCC spike-in controls have been used in previous scRNA-seq studies to identify low quality single cell samples, infer the absolute total number of molecules per cell, and model the technical variability across cells [11,12,14,15]. In our experience, the ERCC controls are not particularly well-suited for any one of these tasks, much less all three. With respect to identifying low quality samples, we indeed observed that samples with no visible cell had a higher percentage of reads mapping to the ERCC controls, as expected. However, there was no clear difference between low and high quality samples in the percentage of ERCC reads or molecules, and thus any arbitrarily chosen cutoff would be associated with considerable error (Figure 1E). With respect to inferring the absolute total number of molecules per cell, we observed that the biological covariate of interest (difference between the three YRI individuals), rather than batch, explained a large proportion of the variance in the ERCC counts (Figure S3), and furthermore that the ERCC controls were also affected by the individual-specific effect on the read-to-molecule conversion rate (Figure 3D). Thus ERCC-based corrected estimates of total number of molecules per cell, across technical or biological replicates, are expected to be biased. Because the batch effects associated with the ERCC controls are driven by the biological covariate of interest, they will also impede the modeling of the technical variation in single cell experiments that confound batch and the biological source of the single cells.

More generally, it is inherently difficult to model unknown sources of technical variation using so few genes [30] (only approximately half of the 92 ERCC controls are detected in typical single cell experiments), and the ERCC controls are also strongly impacted by technical sources of variation even in bulk RNA-seq experiments [31]. Lastly, from a theoretical perspective, the ERCC controls have shorter polyA tails and are overall shorter than mammalian mRNAs. For these reasons, we caution against the reliance of ERCC controls in scRNA-seq studies and highlight that an alternative set of controls that more faithfully mimics mammalian mRNAs and provides more detectable spike-in genes is desired. Our recommendation is to include total RNA from a distant species, for example using RNA from *Drosophila melanogaster* in studies of single cells from humans.

### Outlook

Single cell experiments are ideally suited to study gene regulatory noise and robustness [32,33]. Yet, in order to study the biological noise in gene expression levels, it is imperative that one should be able to effectively estimate and account for the technical noise in single cell gene expression data. Our results indicate that previous single cells gene expression studies may not have been able to distinguish between the technical and the biological components of variation, because single cell samples from each biological condition were processed on a single C1 batch. When technical noise is properly accounted for, even in this small pilot study, our findings indicate pervasive inter-individual differences in gene regulatory noise, independently of the overall gene expression level.

## Methods

### Cell culture of iPSCs

Undifferentiated feeder-free iPSCs reprogrammed from LCLs of Yoruba individuals in Ibadan, Nigeria (abbreviation: YRI) [28] were grown in E8 medium (Life Technologies) [34] on Matrigel-coated tissue culture plates with daily media feeding at 37 °C with 5% (vol/vol) CO2. For standard maintenance, cells were split every 3-4 days using cell release solution (0.5 mM EDTA and NaCl in PBS) at the confluence of roughly 80%. For the single cell suspension, iPSCs were individualized by Accutase Cell Detachment Solution (BD) for 5-7 minutes at 37 °C and washed twice with E8 media immediately before each experiment. Cell viability and cell counts were then measured by the Automated Cell Counter (Bio-Rad) to generate resuspension densities of 2.5 × 105 cells/mL in E8 medium for C1 cell capture.

### Single cell capture and library preparation

Single cell loading and capture were performed following the Fluidigm protocol (PN 1007168). Briefly, 30 *μ*l of C1 Suspension Reagent was added to a 70-*μ*l aliquot of ~17,500 cells. Five *μ*l of this cell mix were loaded onto 10-17 *μ*m C1 Single-Cell Auto Prep IFC microfluidic chip (Fluidigm), and the chip was then processed on a C1 instrument using the cell-loading script according to the manufacturer’s instructions. Using the standard staining script, the iPSCs were stained with StainAlive TRA-1-60 Antibody (Stemgent, PN 09-0068). The capture efficiency and TRA-1-60 staining were then inspected using the EVOS FLCell Imaging System (Thermo Fisher) (Table S1A).

Immediately after imaging, reverse transcription and cDNA amplification were performed in the C1 system using the SMARTer PCR cDNA Synthesis kit (Clontech) and the Advantage 2 PCR kit (Clontech) according to the instructions in the Fluidigm user manual with minor changes to incorporate UMI labeling [20]. Specifically, the reverse transcription primer and the 1:50,000 Ambion^®^ ERCC Spike-In Mix1 (Life Technologies) were added to the lysis buffer, and the template-switching RNA oligos which contain the UMI (5-bp random sequence) were included in the reverse transcription mix [20,35,36]. When the run finished, full-length, amplified, single-cell cDNA libraries were harvested in a total of approximately 13 *μ*l C1 Harvesting Reagent and quantified using the DNA High Sensitivity LabChip (Caliper). The average yield of samples per C1 plate ranged from 1.26-1.88 ng per microliter (Table S1A). A bulk sample, a 40 *μ*l aliquot of ~10,000 cells, was collected in parallel with each C1 chip using the same reaction mixes following the C1 protocol (PN 100-7168, Appendix A).

For sequencing library preparation, tagmentation and isolation of 5’ fragments were performed according to the UMI protocol [20]. Instead of using commercially available Tn5 transposase, Tn5 protein stock was freshly purified in house using the IMPACT system (pTXB1, NEB) following the protocol previously described [37]. The activity of Tn5 was tested and shown to be comparable with the EZ-Tn5-Transposase (Epicentre). Importantly, all the libraries in this study were generated using the same batch of Tn5 protein purification. For each of the bulk samples, two libraries were generated using two different indices in order to get sufficient material for sequencing. All 18 bulk libraries were then pooled and labeled as the "bulk" for sequencing.

### Illumina high-throughput sequencing

The scRNA-seq libraries generated from the 96 single cell samples of each C1 chip were pooled and then sequenced in three lanes on an Illumina HiSeq 2500 instrument using the PCR primer (C1-P1-PCR-2: Bio-GAATGATACGGCGACCACCGAT) as the read 1 primer and the Tn5 adapter (C1-Tn5-U: PHO-CTGTCTCTTATACACATCTGACGC) as the index read primer following the UMI protocol [20].

The master mixes, one mix with all the bulk samples and nine mixes corresponding to the three replicates for the three individuals, were sequenced across four flowcells using a design aimed to minimize the introduction of technical batch effects (Table S1B). Single-end 100 bp reads were generated along with 8-bp index reads corresponding to the cell-specific barcodes. We did not observe any obvious technical effects due to sequencing lane or flow cell that confounded the inter-individual and inter-replicate comparisons.

### Read mapping

To assess read quality, we ran FastQC (http://www.bioinformatics.babraham.ac.uk/projects/fastqc) and observed a decrease in base quality at the 3’ end of the reads. Thus we removed low quality bases from the 3’ end using sickle with default settings [38]. To handle the UMI sequences at the 5’ end of each read, we used umitools [39] to find all reads with a UMI of the pattern NNNNNGGG (reads without UMIs were discarded). We then mapped reads to human genome hg19 (only including chromosomes 1-22, X, and Y, plus the ERCC sequences) with Subjunc [40], discarding non-uniquely mapped reads (option-u). To obtain gene-level counts, we assigned reads to protein-coding genes (Ensembl GRCh37 release 82) and the ERCC spike-in genes using featureCounts [41]. Because the UMI protocol maintains strand information, we required that reads map to a gene in the correct orientation (featureCounts flag -s 1).

In addition to read counts, we utilized the UMI information to obtain molecule counts for the single cell samples. We did not count molecules for the bulk samples because this would violate the assumptions of the UMI protocol, as bulk samples contain far too many unique molecules for the 1,024 UMIs to properly tag them all. First, we combined all reads for a given single cell using samtools [42]. Next, we converted read counts to molecule counts using UMI-tools [43]. UMI-tools counts the number of UMIs at each read start position. Furthermore, it accounts for sequencing errors in the UMIs introduced during the PCR amplification or sequencing steps using a "directional adjacency" method. Briefly, all UMIs at a given read start position are connected in a network using an edit distance of one base pair. However, edges between nodes (the UMIs) are only formed if the nodes have less than a 2x difference in reads. The node with the highest number of reads is counted as a unique molecule, and then it and all connected nodes are removed from the network. This is repeated until all nodes have been counted or removed.

### Filtering cells and genes

We performed multiple quality control analyses to detect and remove data from low quality cells. In an initial analysis investigating the percentage of reads mapping to the ERCC spike-in controls, we observed that replicate 2 of individual NA19098 was a clear outlier (Figure S1A-B). It appeared that too much ERCC spike-in mix was added to this batch, which violated the assumption that the same amount of ERCC molecules was added to each cell. Thus, we removed this batch from all of our analyses.

Next, we kept data from high quality single cells that passed the following criteria:

- Only one cell observed per well
- At least 1,556,255 mapped reads
- Less than 36.4% unmapped reads
- Less than 3.2% ERCC reads
- More than 6,788 genes with at least one read

We chose the above criteria based on the distribution of these metrics in the empty wells (the cutoff is the 95th percentile, Figure S1D-F). In addition, we observed that some wells classified as containing only one cell were clustered with multi-cell wells when plotting 1) the number of gene molecules versus the concentration of the samples (Figure S2A-B), and 2) the read to molecule conversion efficiency (total molecule number divided by total read number) of endogenous genes versus that of ERCC (Figure S2C-D). We therefore established filtering criteria for these misidentified single-cell wells using linear discriminant analysis (LDA). Specifically, LDA was performed to classify wells into empty, one-cell, and two-cell using the discriminant functions of 1) sample concentration and the number of gene molecules (Figure S2A-B) and 2) endogenous and ERCC gene read to molecule conversion efficiency (Figure S2C-D). After filtering, we maintained 564 high quality single cells (NA19098: 142, NA19101: 201, NA19239: 221).

The quality control analyses were performed using all protein-coding genes (Ensembl GRCh37 release 82) with at least one observed read. Using the high quality single cells, we further removed genes with low expression levels for downstream analyses. We removed all genes with a mean log2 cpm less than 2, which did not affect the relative differences in the proportion of genes detected across batches (Figure S9). We also removed genes with molecule counts larger than 1,024 for the correction of collision probability. In the end we kept 13,058 endogenous genes and 48 ERCC spike-in genes.

### Calculate the input molecule quantities of ERCC spiked-ins

According to the information provided by Fluidigm, each of the 96 capture chamber received 13.5 nl of lysis buffer, which contain 1:50,000 Ambion^®^ ERCC Spike-In Mix1 (Life Technologies) in our setup. Therefore, our estimation of the total spiked-in molecule number was 16,831 per sample. Since the relative concentrations of the ERCC genes were provided by the manufacturer, we were able to calculate the molecule number of each ERCC gene added to each sample. We observed that the levels of ERCC spike-ins strongly correlated with the input quantities (r = 0.9914, Figure 1G). The capture efficiency, defined as the fraction of total input molecules being successfully detected in each high quality cell, had an average of 6.1%.

### Subsampling

We simulated different sequencing depths by randomly subsampling reads and processing the subsampled data through the same pipeline described above to obtain the number of molecules per gene for each single cell. To assess the impact of sequencing depth and number of single cells, we calculated the following three statistics:

1. The Pearson correlation of the gene expression level estimates from the single cells compared to the bulk samples. For the single cells, we summed the gene counts across all the samples and then calculated the log_2_ cpm of this pseudo-bulk. For the bulk samples, we calculated the log_2_ cpm separately for each of the three replicates and then calculated the mean per gene.
2. The number of genes detected with at least one molecule in at least one cell.
3. The Pearson correlation of the cell-to-cell gene expression variance estimates from the subsampled single cells compared to the variance estimates using the full single cell data set.

Each data point in Figure 2 represents the mean +/− the standard error of the mean (SEM) of 10 random subsamples of cells. We split the genes by expression level into two groups (6,097 genes each) to highlight that most of the improvement with increased sequencing depth and number of cells was driven by the estimates of the lower half of expressed genes. The data shown is for individual NA19239, but the results were consistent for individuals NA19098 and NA19101. Only high quality single cells (Table S2) were included in this analysis.

### A framework for testing individual and batch effects

Individual effect and batch effect between the single cell samples were evaluated in a series of analyses that examine the potential sources of technical variation on gene expression measurements. These analyses took into consideration that in our study design, sources of variation between single cell samples naturally fall into a hierarchy of individuals and C1 batches. In these sample-level analyses, the variation introduced at both the individual-level and the batch-level was modeled in a nested framework that allows random noise between C1 batches within individuals. Specifically, for each cell sample in individual *i*, replicate *j* and well *k*, we used *y_ijk_* to denote some sample measurement (e.g. total molecule-counts) and fit a linear mixed model with the fixed effect of individual *α_i_* and the random effect of batch *b_ij_*:

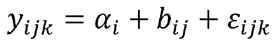

where the random effect *b*_*ij*_-of batch follows a normal distribution with mean zero and variance 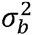 and *ε_ijk_* describes residual variation in the sample measurement. To test the statistical significance of individual effect (i.e., null hypothesis *α*_1_ = *α*_2_ = *α*_3_), we performed a ikelihood ratio test (LRT) to compare the above full model and the reduced model that excludes *α_i_*. To test if there was a batch effect (i.e., null hypothesis 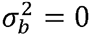), we performed an F-test to compare the variance that is explained by the above full model and the variance due to the reduced model that excludes *b_ij_*.

The nested framework was applied to test the individual and batch effects between samples in the following cases. The data includes samples after quality control and filtering.

1. Total molecule count (on the log_2_ scale) was modeled as a function of individual effect and batch effect, separately for the ERCC spike-ins and for the endogenous genes.
2. Read-to-molecule conversion efficiency was modeled as a function of individual effect and batch effect, separately for the ERCC spike-ins and for the endogenous genes.

### Estimating variance components for per-gene expression levels

To assess the relative contributions of individual and technical variation, we analyzed per-gene expression profiles and computed variance component estimates for the effects of ndividual and C1 batch (Figure S3). The goal here was to quantify the proportion of cell-to-cell variance due to individual (biological) effect and to C1 batch (technical) at the per-gene level. Note that the goal here was different from that of the previous section, where we simply tested for the existence of individual and batch effects at the sample level by rejecting the null hypothesis of no such effects. In contrast, here we fit a linear mixed model per gene where the dependent variable was the gene expression level (log_2_ counts per million) and the independent variables were individual and batch, both modeled as random effects.

The variance parameters of individual effect and batch effect were estimated using a maximum penalized likelihood approach [44], which can effectively avoid the common issue of zero variance estimates due to small sample sizes (there were three individuals and eight batches). We used the blmer function in the R package blme and set the penalty function to be the logarithm of a gamma density with shape parameter = 2 and rate parameter tending to zero.

The estimated variance components were used to compute the sum of squared deviations for individual and batch effects. The proportion of variance due to each effect is equal to the relative contribution of the sum of squared deviations for each effect compared to the total sum of squared deviations per gene. Finally, we compared the estimated proportions of variance due to the individual effect and the batch effect, across genes, using a non-parametric one-way analysis of variance (Kruskal-Wallis rank sum test).

### Normalization

We transformed the single cell molecule counts in multiple steps (Figure 4). First, we corrected for the collision probability using a method similar to that developed by Grün et al. [12]. Essentially we corrected for the fact that we did not observe all the molecules originally in the cell. The main difference between our approach and that of Grün et al. [12] was that we applied the correction at the level of gene counts and not individual molecule counts. Second, we standardized the molecule counts to log2 counts per million (cpm). This standardization was performed using only the endogenous gene molecules and not the ERCC molecules. Third, we corrected for cell-to-cell technical noise using the ERCC spike-in controls. For each single cell, we fit a Poisson generalized linear model (GLM) with the log_2_ expected ERCC molecule counts as the independent variable, and the observed ERCC molecule counts as the dependent variable, using the standard log link function. Next we used the slope and intercept of the Poisson GLM regression line to transform the log2 cpm for the endogenous genes in that cell. This is analogous to the standard curves used for qPCR measurements, but taking into account that lower concentration ERCC genes will have higher variance from Poisson sampling. Fourth, we removed technical noise between the eight batches (three replicates each for NA19101 and NA19239 and two replicates for NA19098). We fit a linear mixed model with a fixed effect for individual and a random effect for the eight batches and removed the variation captured by the random effect (see the next section for a detailed explanation).

For the bulk samples, we used read counts even though the reads contained UMIs. Because these samples contained RNA molecules from ~10,000 cells, we could not assume that the 1,024 UMIs were sufficient for tagging such a large number of molecules. We standardized the read counts to log_2_ cpm.

### Removal of technical batch effects

Our last normalization step adjusted the transformed log_2_ gene expression levels for cell-to-cell correlation within each C1 plate. The algorithm mimics a method that was initially developed for adjusting within-replicate correlation in microarray data [24]. We assumed that for each gene *g*, cells that belong to the same batch *j* are correlated, for batches *j* = 1,…,8. The batch effect is specific to each C1 plate and is independent of biological variation across individuals.

We fit a linear mixed model for each gene *g* that includes a fixed effect of individual and a random effect for within-batch variation attributed to cell-to-cell correlation in each C1 plate:

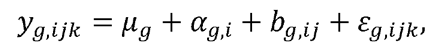

where *y_g,ijk_* denotes log_2_ counts-per-million (cpm) of gene *g* in individual *i*, replicate *j*, ind cell *k*; *i* = *NA*19098,*NA*19101,*NA*19239, *j* = l,…,*n_i_* with *n_i_* the number of replicates in ndividual *i*, *k* = 1,…,*n_ij_* with *n_ij_* the number of cells in individual *i* replicate *j*. *μ_g_* denotes the mean gene expression level across cells, *α_g,i_* quantifies the individual effect on mean gene expression, *b_g,ij_* models the replicate effect on mean expression level (assumed to be tochastic, independent, and identically distributed with mean 0 and variance 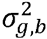). Finally, *ε_g,ijk_* escribes the residual variation in gene expression.

Batch-corrected expression levels were computed as

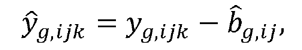

where 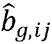 are the least-squares estimates. The computations in this step were done with the gls.series function of the limma package [45].

### Measurement of gene expression noise

While examining gene expression noise (using the coefficient of variation or CV) as a function of mean RNA abundance across C1 replicates, we found that the CV of molecule counts among endogenous genes and ERCC spike-in controls suggested similar expression variability patterns. Both endogenous and ERCC spike-in control CV patterns approximately followed an over-dispersed Poisson distribution (Figure S10A-C), which is consistent with previous studies [11,20]. We computed a measure of gene expression noise that is independent of RNA abundance across individuals [46,47]. First, squared coefficients of variation (CVs) for each gene were computed for each individual and also across individuals, using the batch-corrected molecule data. Then we computed the distance of individual-specific CVs to the rolling median of global CVs among genes that have similar RNA abundance levels. These transformed individual CV values were used as our measure of gene expression noise. Specifically, we computed the adjusted CV values as follows:

1. Compute squared CVs of molecule counts in each individual and across individuals.
2. Order genes by the global average molecule counts.
3. Starting from the genes with the lowest global average gene expression level, for every sliding window of 50 genes, subtract log_10_ median squared CVs from log_10_ squared CVs of each cell line, and set 25 overlapping genes between windows. The computation was performed with the rollapply function of the R zoo package [48]. After this transformation step, CV no longer had a polynomial relationship with mean gene molecule count (Figure S10D-E).

### Identification of genes associated with inter-individual differences in regulatory noise

To identify differential noise genes across individuals, we computed median absolute deviation (MAD)-a robust and distribution-free dissimilarity measure for gene *g*:

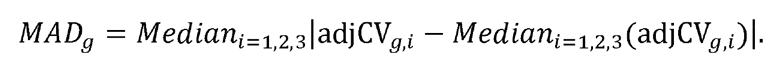

Large values of *MAD_g_* suggest a large deviation from the median of the adjusted CV values. We identified genes with significant inter-individual differences using a permutation-based approach. Specifically, for each gene, we computed empirical *P*-values based on 300,000 permutations. In each permutation, the sample of origin labels were shuffled between cells. Because the number of permutations in our analysis was smaller than the maximum possible number of permutations, we computed the empirical *P*-values as 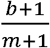 where *b* is the
number of permuted MAD values greater than the observed MAD value, and *m* is the number of permutations. Adding 1 to *b* avoided an empirical *P*-value of zero [49].

### Gene enrichment analysis

We used ConsensusPATHDB [50] to identify GO terms that are over-represented for genes whose variation in single cell expression levels were significantly difference between individuals.

### Individual assignment based on scRNA-seq reads

We were able to successfully determine the correct identity of each single cell sample by examining the SNPs present in their RNA sequencing reads. Specifically, we used the method verifyBamID (https://github.com/statgen/verifyBamID) developed by Jun et al., 2012 [27], which detects sample contamination and/or mislabeling by comparing the polymorphisms observed in the sequencing reads for a sample to the genotypes of all individuals in a study. For our test, we included the genotypes for all 120 Yoruba individuals that are included in the International HapMap Project [28]. The genotypes included the HapMap SNPs with the 1000 Genomes Project SNPs [51] imputed, as previously described [52]. We subset to include only the 528,289 SNPs that overlap Ensembl protein-coding genes. verifyBamID used only 311,848 SNPs which passed its default thresholds (greater than 1% minor allele frequency and greater than 50% call rate). Using the option --best to return the best matching individual, we obtained 100% accuracy identifying the single cells of all three individuals (Figure S8B).

### Data and code availability

The data have been deposited in NCBI’s Gene Expression Omnibus [53] and are accessible through GEO Series accession number GSE77288 (http://www.ncbi.nlm.nih.gov/geo/query/acc.cgi?acc=GSE77288). The code and processed data are available at https://github.com/jdblischak/singleCellSeq. The results of our analyses are viewable at https://jdblischak.github.io/singleCellSeq/analysis.

## Acknowledgment

We thank members of the Pritchard, Gilad, and Stephens laboratories for valuable discussions during the preparation of this manuscript. This work was funded by NIH grant HL092206 to YG and HHMI funds to JKP. PYT is supported by NIH T32HL007381.

## Supplemental Figures

**Figure S1.**
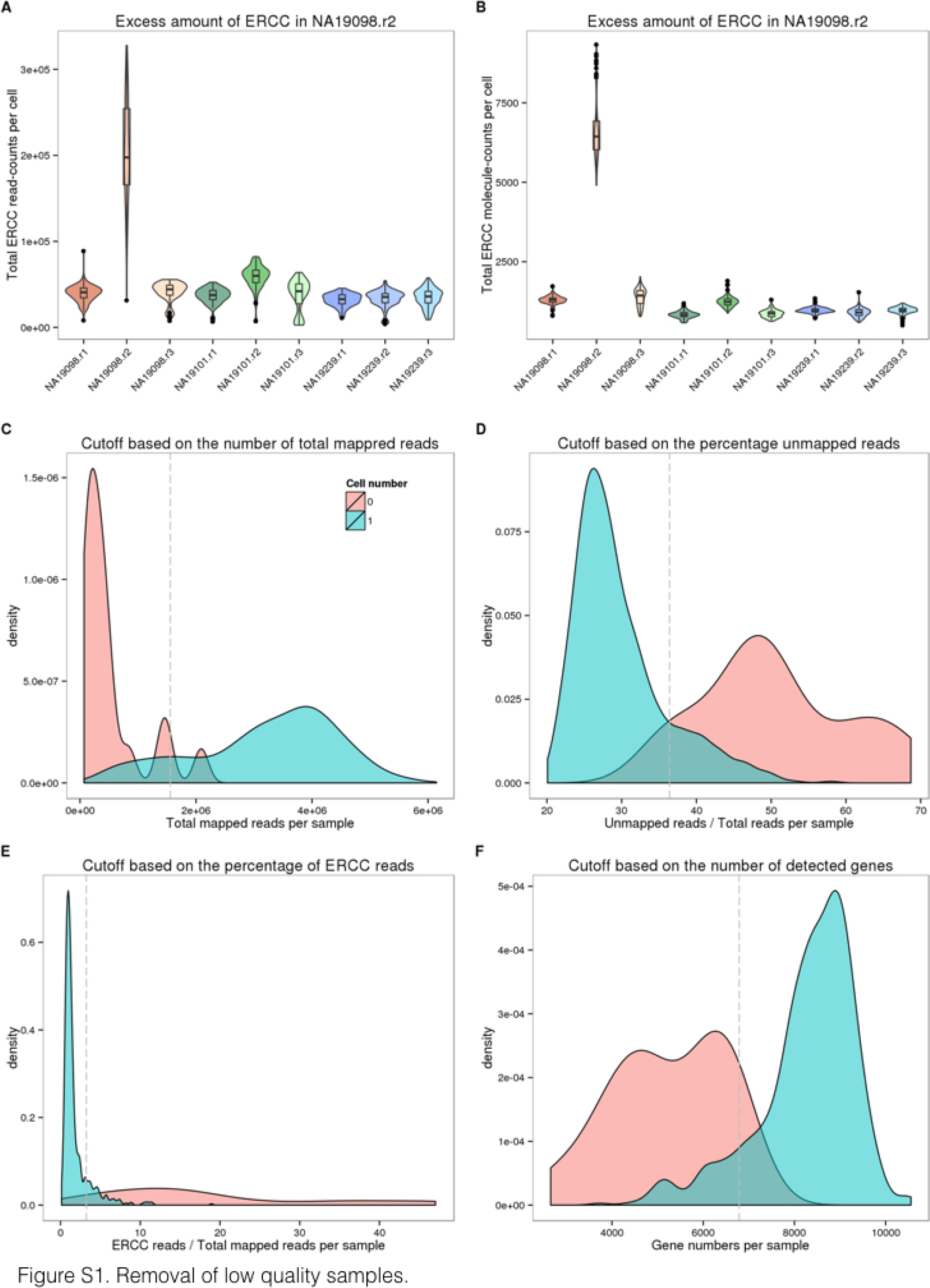
Removal of low quality samples. Violin plots of the total read-counts of ERCC spike-in controls in (A) and the total molecule-counts in (B) in single cell samples. The three colors represent the three individuals (NA19098 in red, NA19101 in green, and NA19239 in blue). (C-F) Density plots of the distributions of the total mapped reads in (C), the percentage of unmapped reads in (D), the percentage of ERCC reads in (E), and the number of detected genes in (F). The dash lines indicate the cutoffs based on the 95th percentile of the samples with no cells.

**Figure S2.**
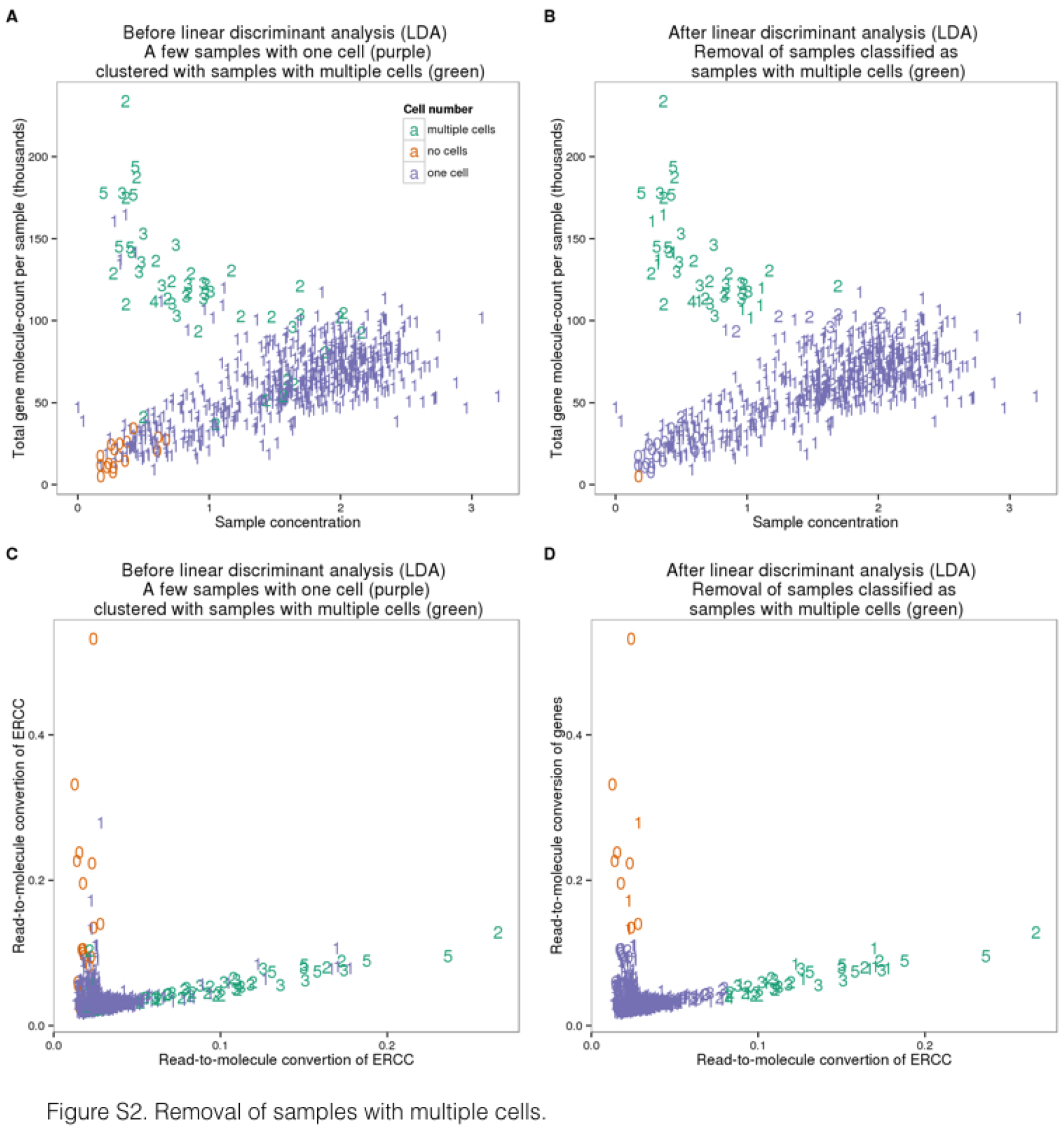
Removal of samples with multiple cells. Scatterplots of the three groups of samples (no cell in green, single-cell in orange, and two or more cells in purple) before (A) and after (B) the linear discriminant analysis (LDA) using sample concentration of cDNA amplicons (ng/*μ*l) and the number of detected genes. (C and D) Similarly, LDA was performed to identify potential multi-cell samples using the read-to-molecule conversion efficiency (total molecule-counts divided by total read-counts per sample) of endogenous genes and ERCC spike-in controls. Scatterplots of before and after the LDA in (C) and (D), respectively. The numbers indicate the number of cells observed in each cell capture site.

**Figure S3.**
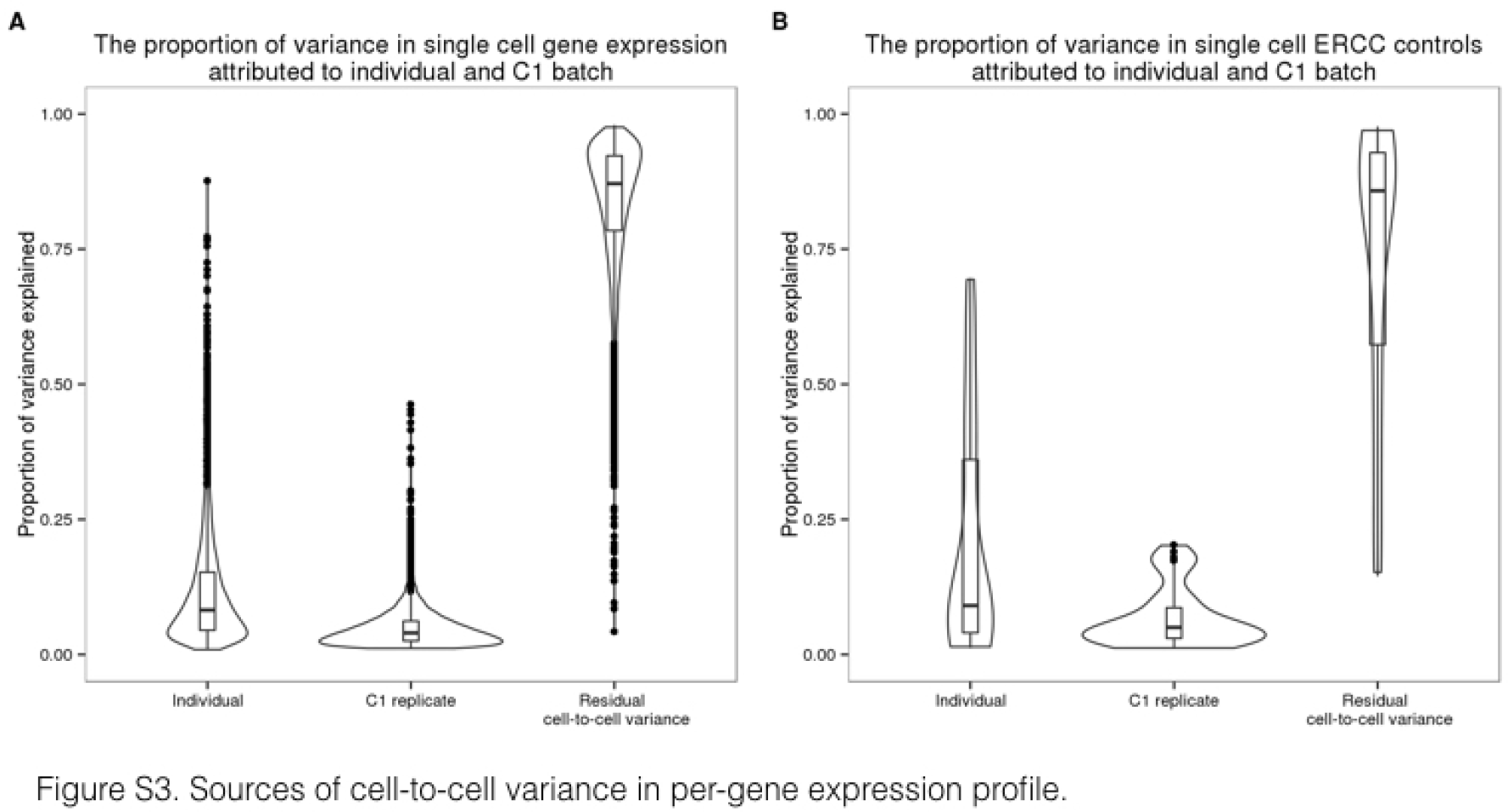
Sources of cell-to-cell variance in per-gene expression profile. Violin plots of the proportion of per-gene cell-to-cell variance that was due to individual sample of origin, different C1 replicates, and other single cell sample differences. These results were calculated from the molecule counts before normalization and batch correction. Endogenous genes are shown in (A) and the ERCC spike-in controls in (B).

**Figure S4.**
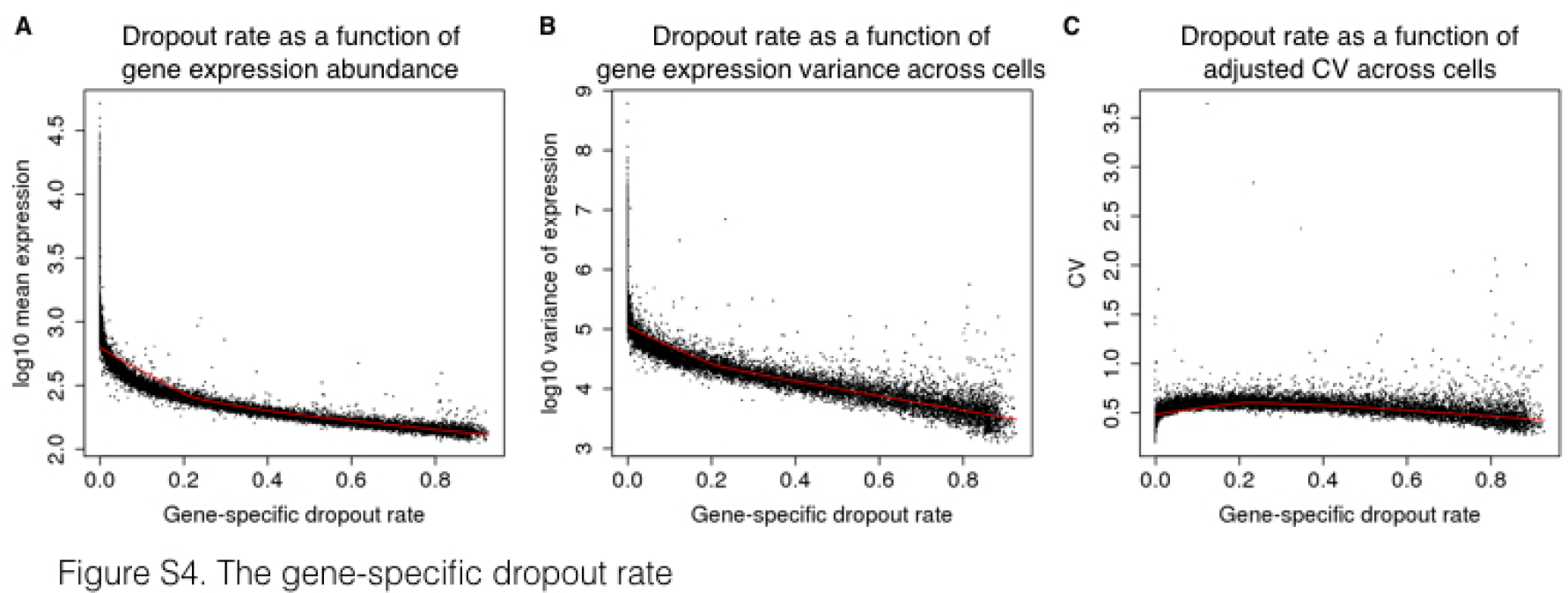
The gene-specific dropout rate. The gene-specific dropout rate (the proportion of cells in which the gene is undetected) and its relationship with log_10_ mean expression in (A), with log_10_ variance of expression in (B), and with the CV in (C) of the cells in which the gene is expressed (cells in which at least one molecule of the given gene was detected). Each point represents a gene, and red lines indicate the predicted values using locally weighted scatterplot smoothing (LOESS).

**Figure S5.**
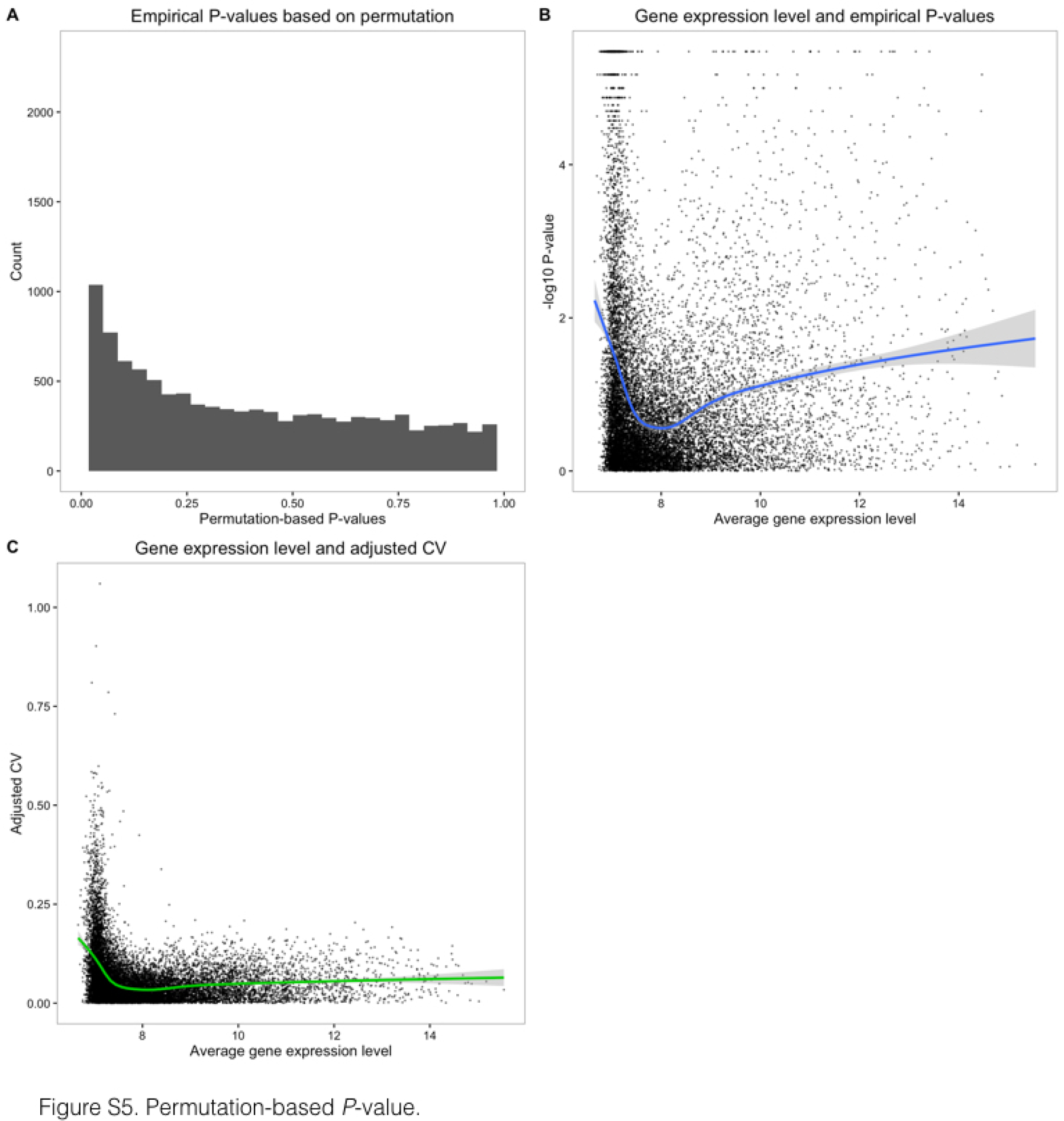
Permutation-based P-value. (A) Histogram of empirical P-values based on 300,000 permutations. (B) −log_10_ empirical P-values are plotted against average gene expression levels. Blue line indicates the fitted relationship between −log_10_ *P*-values and average log_2_ gene expression levels of cells that were detected as expressed, using locally weighted scatterplot smoothing (LOESS). (C) Median of Absolute Deviation (MAD) of genes versus average gene expression levels. Green line indicates the fitted relationship (LOESS) between the MAD values and average log_2_ gene expression levels of cells in which the gene was detected as expressed.

**Figure S6.**
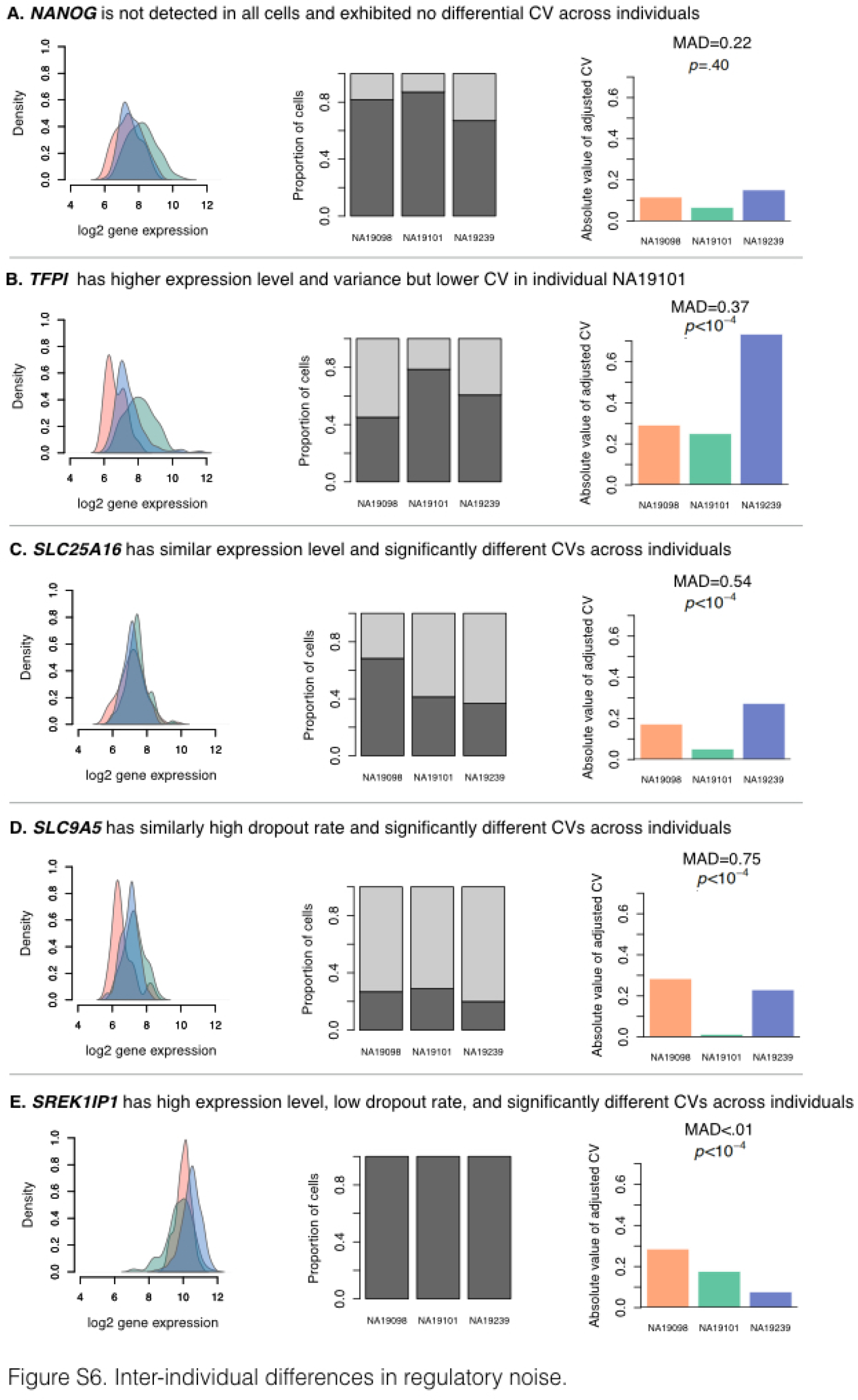
Inter-individual differences in regulatory noise. These 5 example genes illustrate various patterns of cell-to-cell gene expression variance. For each gene, the left panel shows the distribution of the log_2_ gene expression levels (considering only cells in which the gene is detected as expressed), the middle panel shows the proportion of cells in which the gene is detected as expressed (dark grey) and the dropout rate (light grey) for each individual, and the right panel shows the absolute value of adjusted CV for each individual, along with the corresponding gene-specific MAD (median of absolute deviation) value and P-value. The three colors in the upper and lower panel represent the individuals (NA19098 in red, NA19101 in green, and NA19239 in blue).

**Figure S7.**
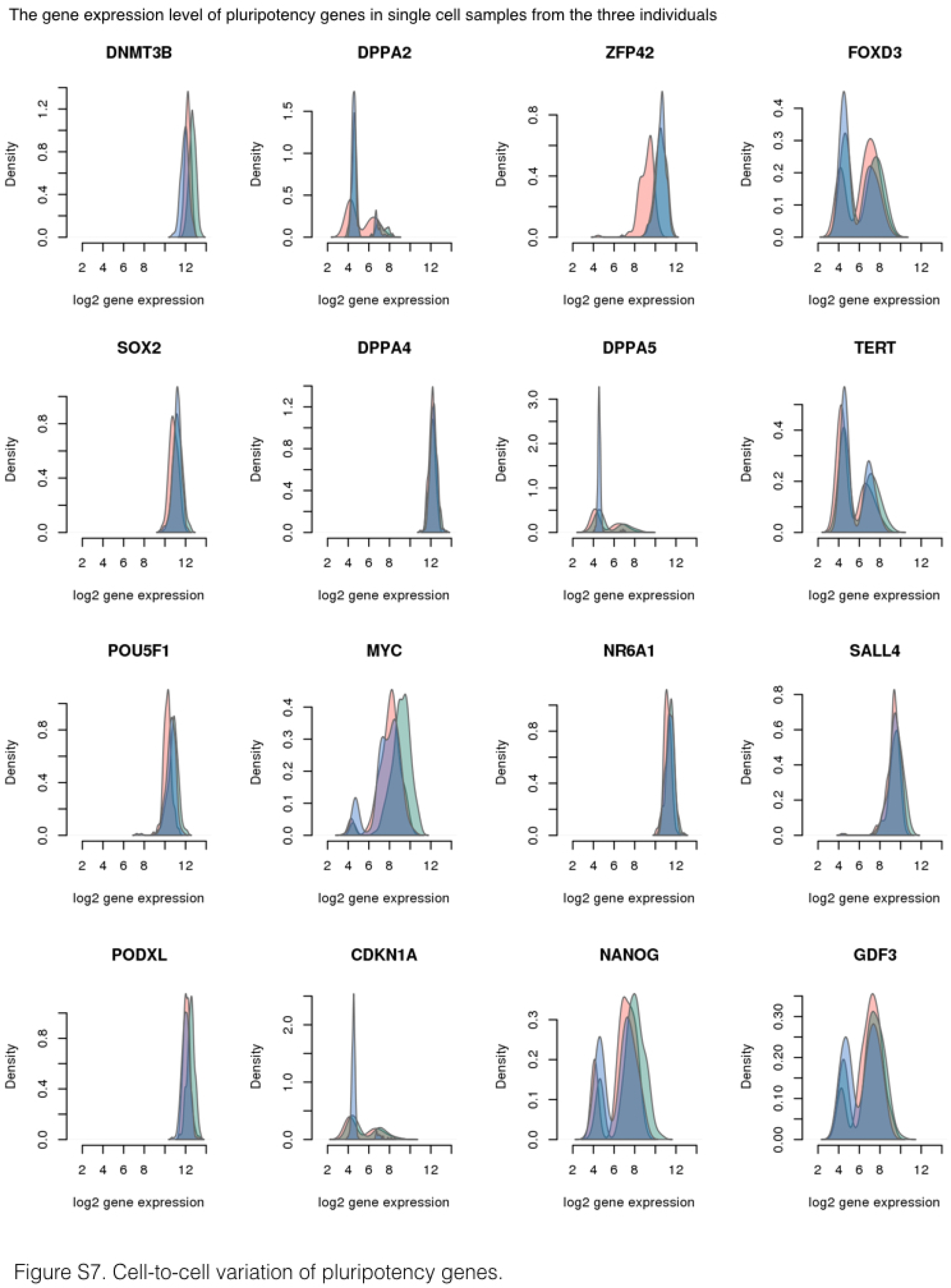
Cell-to-cell variation of pluripotency genes. Density plots of the distribution of log_2_ gene expression of key pluripotency genes across all single cells by individual. The peaks with lower gene expression values (log2 around 4) represent the cells in which the gene is undetected. The three colors represent the three individuals (NA19098 is in red, NA19101 in green, and NA19239 in blue).

**Figure S8.**
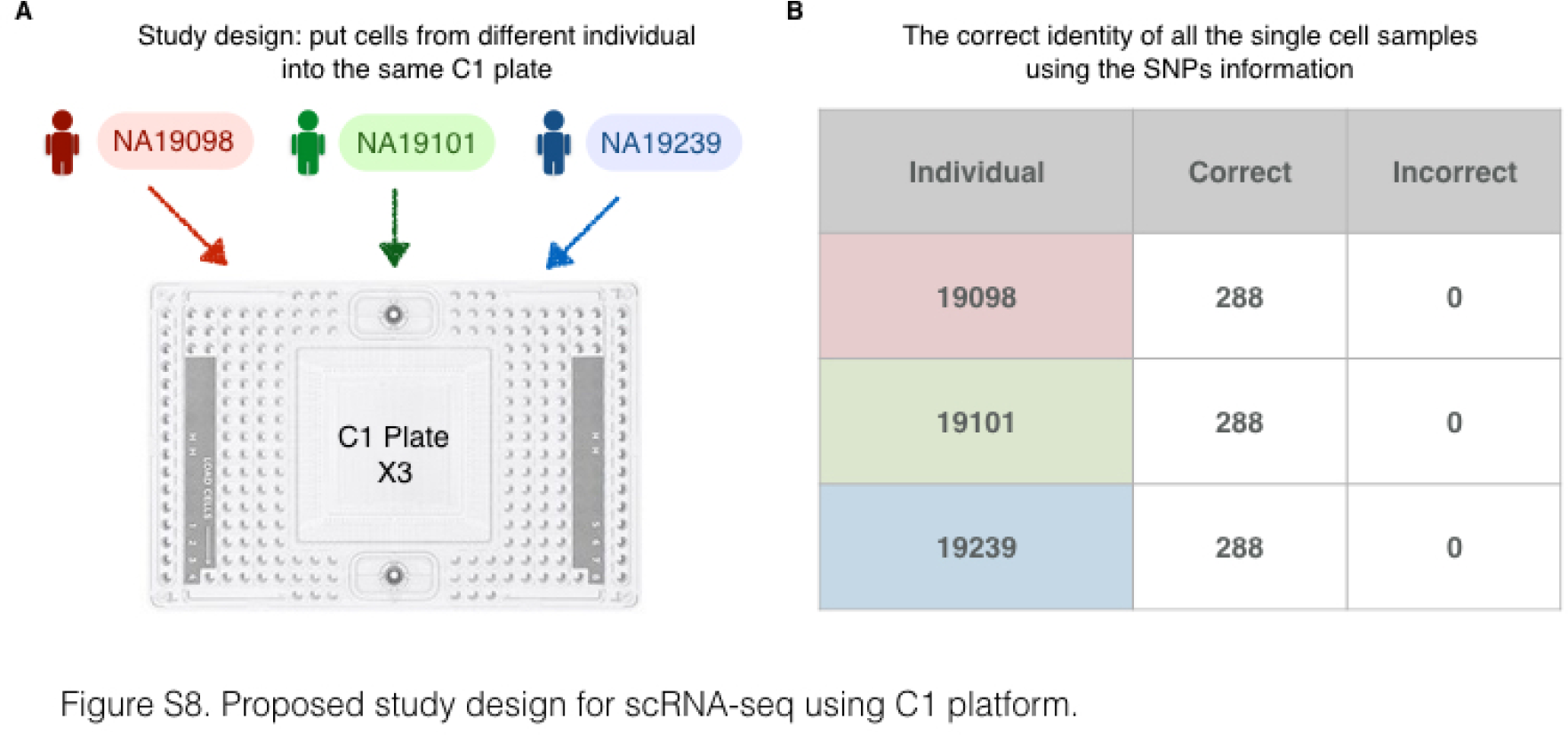
Proposed study design for scRNA-seq using C1 platform. (A) A balanced study design consisting of multiple individuals within a C1 plate and multiple C1 replicates to fully capture the batch effect across C1 plates and further retrieve the maximum amount of biological information. (B) The correct identity of each single cell sample was determined by examining the SNPs present in their RNA sequencing reads.

**Figure S9.**
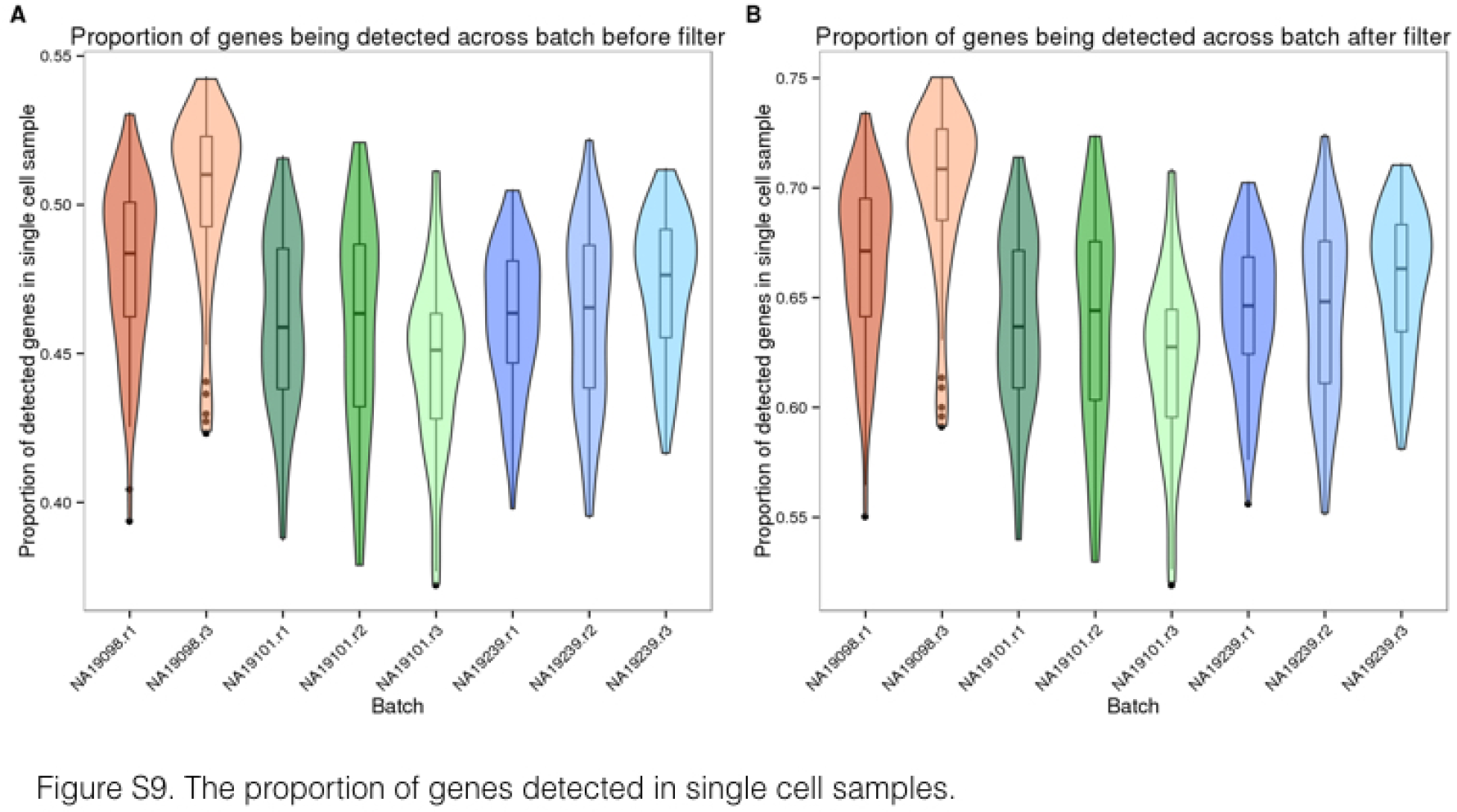
The proportion of genes detected in single cell samples. Violin plots of the proportion of genes detected, computed by the total number of detected genes in each single cell divided by the total number of genes detected across all single cells, before in (A) and after in (B) the removal of genes with low expression. The three colors represent the three individuals (NA19098 is in red, NA19101 in green, and NA19239 in blue).

**Figure S10.**
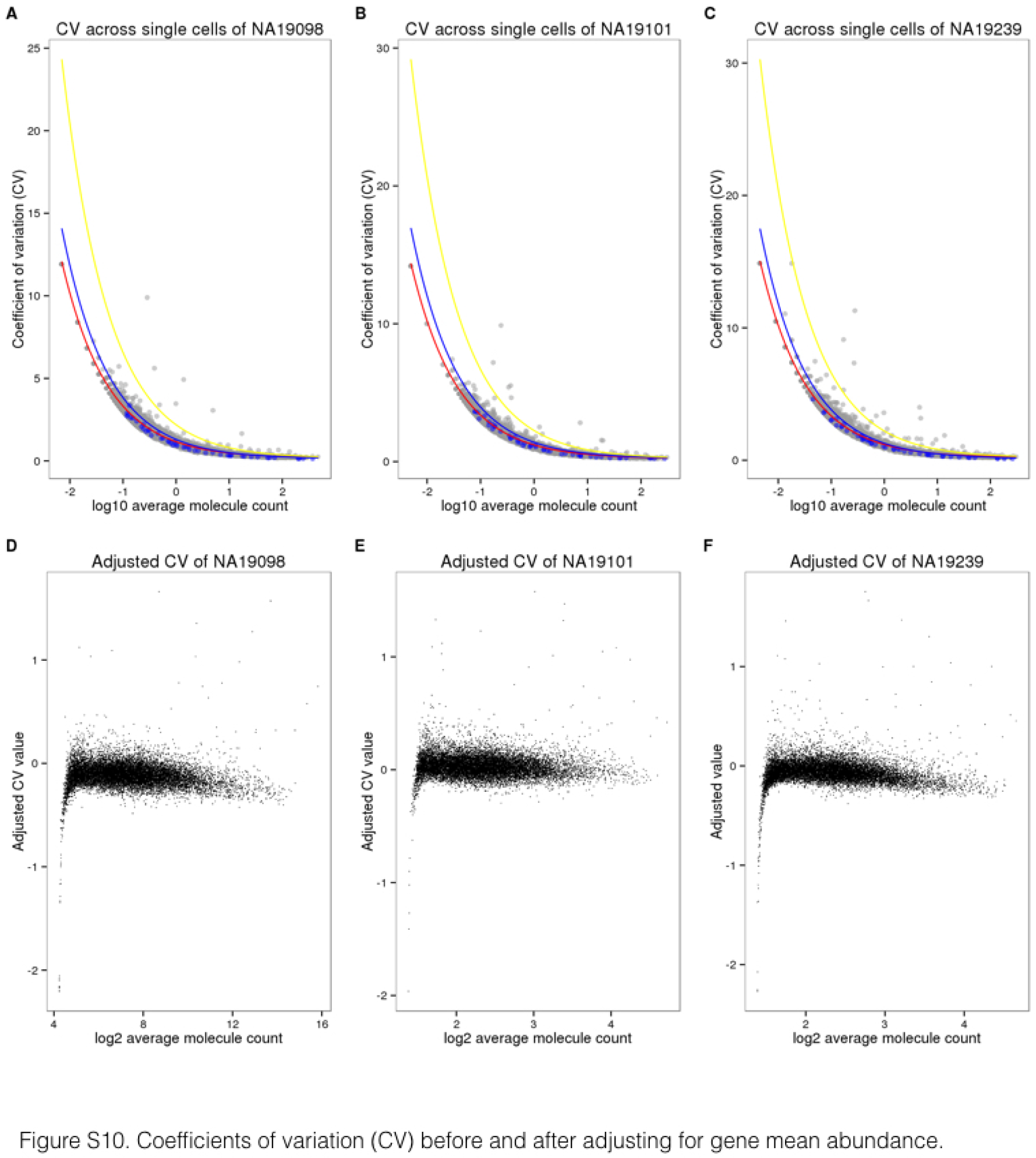
Coefficients of variation (CV) before and after adjusting for gene mean abundance. (A-C) CV plotted against average molecule counts across all cells for each individual [20]. Grey points represent endogenous genes, and blue points represent ERCC spike-in controls. The curves indicate the expected CV under three different scenarios. Red curve depicts the expected CV of the endogenous genes while assuming a Poisson distribution with no over-dispersion. Likewise, blue curve depicts the expected CVs of the ERCC spike-in controls under the Poisson assumption. Yellow curve depicts the expected CVs of an overdispersed Poisson distribution for which standard deviation is three times the ERCC spike-in controls. (D-F) Adjusted CV values of each gene including all cells are plotted against log™ of the average molecule counts for each individual.

### Supplemental Tables

**Table S1.** Data collection. (A) iPSCs were sorted using the 10-17 *μ*m IFC plates with the staining of the pluripotency marker, TRA1-60. Single cell occupancy is the percentage of occupied capture sites containing one single cell. The average cDNA concentration was measured by the HT DNA high sensitivity LabChip (Caliper). (B) The 96 single cell libraries from one C1 plate were pooled and sequenced in three HiSeq lanes. The pooled samples were assigned across the four 8-lane flowcells.

| Cell line | Passage | Input viability | No cell | Multiple cells | % single cell occupancy | TRA1-60 negative | % TRA1-60 | Date | Ave cDNA con (ng/ul) | Replicate for seq |
| --- | --- | --- | --- | --- | --- | --- | --- | --- | --- | --- |
| 19098 | 12+15 | 89% | 2 | 2 | 95.83 | 2 | 97.92 | 11/07/2014 | N/A | (1) |
| 19098 | 12+18 | 90% | 2 | 4 | 93.75 | 0 | 100.00 | 11/14/2014 | 1.88 | (2) |
| 19098 | 12+20 | 83% | 3 | 10 | 86.46 | 0 | 100.00 | 11/22/2014 | 1.40 | (3) |
| 19101 | 12+16 | 70% | 2 | 3 | 94.79 | 9 | 90.63 | 11/13/2014 | 1.81 | (1) |
| 19101 | 12+19 | 94% | 5 | 3 | 91.67 | 0 | 100.00 | 11/23/2014 | 1.38 | (2) |
| 19101 | 12+19 | 69% | 1 | 19 | 79.17 | 0 | 100.00 | 11/24/2014 | 1.26 | (3) |
| 19239 | 12+16 | 85% | 1 | 2 | 96.88 | 4 | 95.83 | 11/11/2014 | 1.60 | (1) |
| 19239 | 12+18 | 75% | 1 | 6 | 92.71 | 5 | 94.79 | 11/17/2014 | 1.55 | (2) |
| 19239 | 12+19 | 93% | 5 | 7 | 87.50 | 2 | 97.92 | 11/21/2014 | 1.70 | (3) |

Table S1. Data collection.
| Flowcell 1 | Flowcell 2 | Flowcell 3 | Flowcell 4 |
| --- | --- | --- | --- |
| Bulk | 19098 (2) | 19098 (3) | 19239 (1) |
| 19098 (1) | 19239 (3) | 19101 (1) | 19101 (2) |
| 19239 (2) | 19098 (1) | 19098 (2) | 19098 (3) |
| 19101 (3) | 19239 (2) | 19239 (3) | 19101 (1) |
| 19239 (1) | 19101 (3) | Bulk | 19098 (2) |
| 19101 (2) | 19239 (1) | 19098 (1) | 19239 (3) |
| 19098 (3) | 19101 (2) | 19239 (2) | Bulk (all 9) |
| 19101 (1) | Bulk | 19101 (3) |  |

**Table S2. High quality single cell samples.**

List of the 564 high quality single cell samples.

**Table S3. Genes associated with inter-individual differences in regulatory noise.**

List of genes that we classified the estimates of regulatory noise as significantly different across individuals (empirical permutation *P* < 10^−4^). There are a total of 560 genes.

**Table S4. Gene ontology analysis of the genes associated with inter-individual differences in regulatory noise.**

